# ER-export and ARFRP1/AP-1-dependent delivery of SARS-CoV-2 Envelope to lysosomes controls late stages of viral replication

**DOI:** 10.1101/2021.06.30.450614

**Authors:** G.J Pearson, H. Mears, M. Broncel, A.P. Snijders, D.L.V. Bauer, J.G. Carlton

## Abstract

The β-coronavirus SARS-CoV-2 is the causative agent of the global Covid-19 pandemic. Coronaviral Envelope (E) proteins are pentameric viroporins that play essential roles in assembly, release and pathogenesis. We developed an inert tagging strategy for SARS-CoV-2 E and find that it localises to the Golgi and to lysosomes. We identify sequences in E, conserved across *Coronaviridae,* responsible for ER-to-Golgi export, and relate this activity to interaction with COP-II via SEC24. Using proximity biotinylation, we identify host-cell factors that interact with E and identify an ARFRP1/AP-1 dependent pathway allowing Golgi-to-lysosome trafficking of E. We identify sequences in E that bind AP-1, are conserved across β-coronaviruses and allow E to be trafficked from Golgi to lysosomes. We show that E acts to deacidify lysosomes and by developing a *trans*-complementation assay, we show that both lysosomal trafficking of E and its viroporin activity are necessary for efficient viral replication and release.

## INTRODUCTION

Severe Acute Respiratory Syndrome Coronavirus-2 (SARS-CoV-2) is an enveloped, β-coronavirus with a positive-sense RNA genome encoding at least 29 different proteins^1^. Late events in the β-coronaviral lifecycle are orchestrated by 4 of these proteins, the RNA-binding protein Nucleocapsid (N) and the three transmembrane proteins Spike (S), Membrane (M) and Envelope (E). Viral assembly occurs on internal membranes and involves the budding of nascent particles into the secretory pathway lumen^2,3^. The structural proteins M and E are thought to be necessary for viral budding^4,5^ with incorporation of N allowing packaging of the viral genome^6^. At steady state, coronaviral E proteins are known to localise to Golgi membranes, sites of coronaviral particle assembly^3,7,8^. E is predicted to form a pentameric cation channel^9,10^ and is only a minor component of coronavirus virions^11^, suggesting that it plays important roles in manipulating the biology of the host. Indeed, the channel activity of E contributes to Acute Respiratory Distress Syndrome (ARDS)-like pathological damage of E-expressing cells in both cellular and animal models^10^. Recombinant CoVs lacking E exhibit defects in viral maturation and replication. For example, a SARS-CoV that lacks the E gene is attenuated *in-vitro* and *in-vivo*^12^, a recombinant Murine Hepatitis Virus (MHV) lacking E can replicate, but produces smaller plaques *in-vitro*^13^, and a recombinant Transmissible Gastroenteritis Virus (TGEV) lacking E is blocked in viral release with virions retained in the secretory pathway^14^. These data suggest that E controls late events in the coronavirus lifecycle that allow virus production and maturation^3^. Whilst viral egress was assumed to occur via the canonical secretory pathway, recent data suggest that β-coronaviruses can be delivered to deacidified lysosomes for atypical secretion via lysosomal exocytosis^15^. Expression of E has been shown to cause deacidification of lysosomes^16^ and a mutation (E^T9I^) in currently circulating omicron (B.1.1.529) variants that eliminates a polar pore-lining residue compromises lysosomal deacidification and leads to a reduced viral load^17^, which is suggested to contribute to the reduced pathogenicity of this variant. Here, we asked how E was delivered to lysosomes to exert these effects. We identify sequence elements conserved across β-coronaviral E proteins that allow engagement with transport machineries allowing both ER-to-Golgi traffic and Golgi-to-lysosome traffic and we identify the Golgi-localised GTPase ARFRP1 as being essential for the recruitment of AP-1 to the Golgi and for coordinating an AP-1-dependent trafficking route for delivering E to lysosomes. By developing a *trans*-complementation assay for E sub-genomic mRNA, we demonstrate the importance of this trafficking pathway for late stages of the SARS-CoV-2 lifecycle.

## RESULTS

### Internally tagged SARS-CoV-2 Envelope traffics to and deacidifies lysosomes

We generated tagged versions of E to investigate its intracellular trafficking itinerary. We were surprised to find N- or C-terminal HaloTag (HT) fusions restricted the localisation of E to the Endoplasmic Reticulum (ER) (Fig. 1A-1C), suggesting that canonical tagging disrupts the proper localisation of this protein. We found that placement of HT at internal positions either immediately after the transmembrane domain (E-HT^Site3^), or in a region of the cytoplasmic tail (E-HT^Site4^) of E allowed steady state localisation to Golgi membranes (Fig. 1A-1C), consistent with known localisation for E proteins^3^. We confirmed these localisations using mEmerald in place of HT (Fig. S1A and S1B) and devised a *qu*antitative *i*maging-based *l*ocalisation *t*able (*quilt*) to depict E’s position within the secretory pathway (Fig. 1C and Fig. S1B). Unless otherwise indicated, experiments hereafter employ internal tags placed at Site3, with analysis performed after 16-18 hours of expression to limit the toxicity associated with expression of this viroporin (Fig. S1C)^10^.

**Figure 1.**
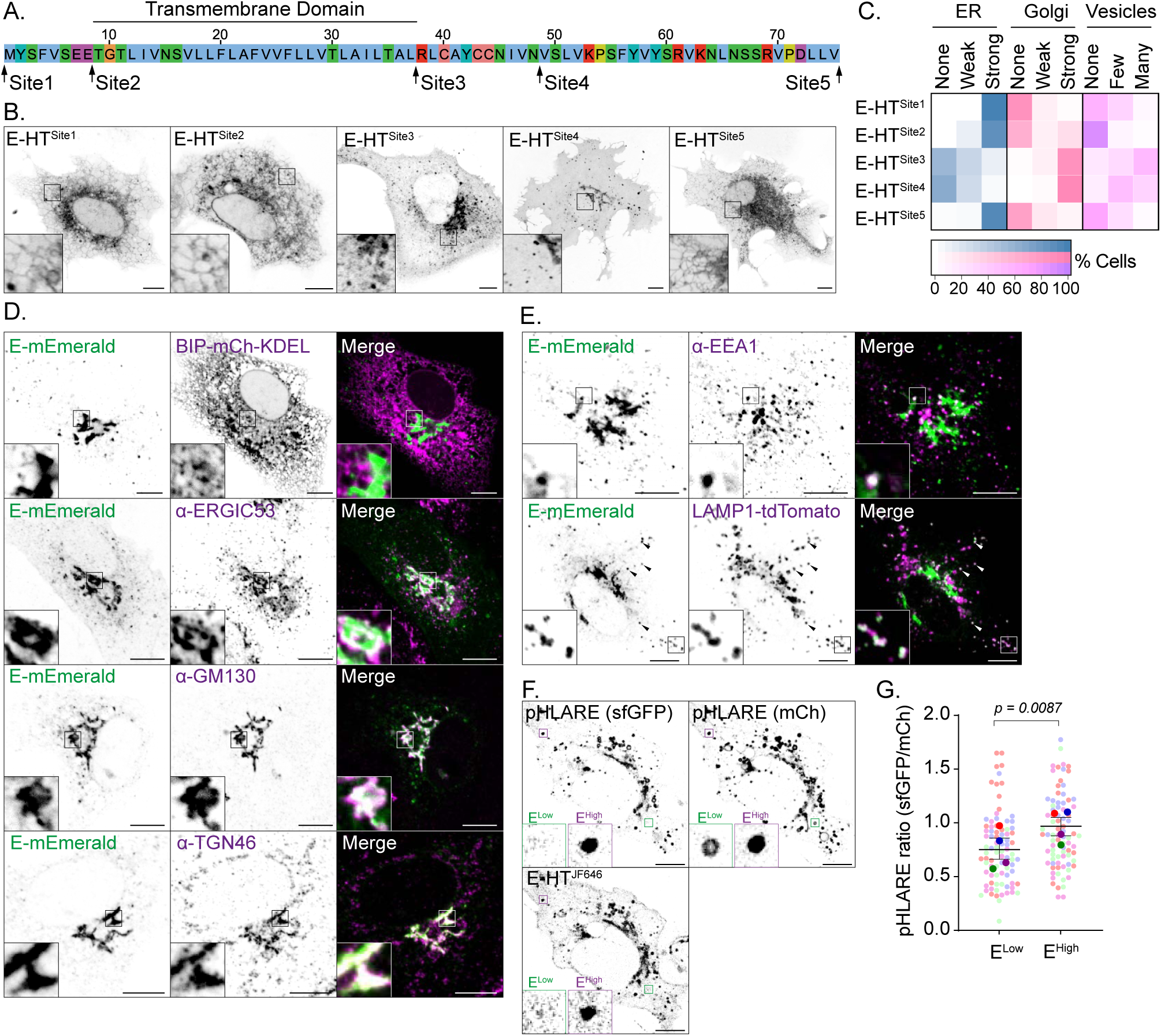
Placement of internal tags allows visualisation of the trafficking itinerary of SARS-CoV-2 Envelope. (**A**) Sequence of Envelope (E) protein from SARS-CoV-2 indicating the position of internal insertion sites. Amino acids coloured according to ClustalX colour criteria (**B**) Representative live images of VeroE6 cells transfected with plasmids encoding the indicated Janelia Fluor (JF) 646-illuminated E-HaloTag (HT) fusion proteins. (**C**) Quantitative Imaging-based Localisation Table (Quilt) of the indicated HT-fusion proteins from 50 imaged cells in B. (**D, E**) Representative images of VeroE6 cells transfected with plasmids encoding E-mEmerald^Site3^ fixed and stained with antisera raised against ERGIC53, GM130, TGN46 or EEA1, or that were co-transfected with plasmids encoding BIP-mCh-KDEL or LAMP1-tdTomato. Images representative of 15 imaged cells in each case. Arrowheads indicate colocalised E-mEmerald and LAMP1-tdTomato. (**F**) Representative image of VeroE6 cells expressing pHLARE and JF646-illuminated E-HT. Examples of E-HT-low and E-HT-high lysosomes are displayed. (**G**) Superplot of ratiometric imaging of sfGFP and mCherry in JF646-high and JF646-low lysosomes within the same cell. Each presented data point represents the mean sfGFP/mCh signal of each mCh-positive lysosome in the cell binned into high or low classes based upon its JF646 signal. Mean ± S.E. displayed, with significance determined by paired 2-tailed T-test. In microscopy panels, scale bars are 10 μm.

We found good colocalization of E-mEmerald with endogenous GM130 and TGN46 (Fig. 1D). We observed no co-localisation with an mCherry targeted to the ER-lumen, and we observed partial colocalization with the Golgi-localised pool of ERGIC53. These data suggest at steady state, E-mEmerald is exported efficiently from the ER and reaches the Golgi. In addition to the predominate perinuclear Golgi localisation, we noticed that E-mEmerald decorated punctate structures in the cytoplasm (Fig. 1B-1E). We found good colocalisation of these peripheral puncta with LAMP1, a marker of late endosomes and lysosomes and occasional colocalization with EEA1 (Fig. 1E), suggesting lysosomal access is gained via endosomes. Both perinuclear and punctate localisation of E-HT was confirmed in cells expressing all 4 SARS-CoV-2 structural proteins, indicating that these localisations are preserved during particle assembly (Fig. S1D).

Coronaviral E proteins assemble into pentameric viroporins^9^. We used Fluorescence Lifetime Imaging-Forster Radius Energy Transfer (FLIM-FRET) to confirm that the lifetime of E-mEmerald in post-Golgi vesicular structures was reduced in the presence of TMR-labelled E-HT, suggesting that E oligomerises in these organelles (Fig. S1E and S1F). We transfected E-HT into VeroE6 cells and, using a recently described tandem-fluorescent reporter or lysosomal pH (pHLARE)^18^ (Fig. S1G and S1H), we found that lysosomes containing higher levels of E-HT were deacidified relative to lysosomes containing lower levels of E-HT (Fig.1F and 1G). These data suggest that whilst E localises predominantly to Golgi membranes, a pool of E is trafficked onwards to lysosomes and allows pH-neutralisation in these organelles.

### Identification of an ER-export motif in SARS-CoV-2 Envelope

Transmembrane proteins are co-translationally inserted into the ER. We next performed alanine-scanning mutagenesis through the cytosolic tail of E to identify sequences required for its trafficking to the Golgi and onwards towards lysosomes (Fig. S2). We found that mutation of the terminal 4 amino acids to Alanine (E-mEmerald^M9^) restricted E to its site of biosynthesis in the ER (Fig. 2A-2D). Grafting these C-terminal amino acids onto E-HT^Site5^ restored anterograde traffic of this protein (Fig. 2E - Fig. 2G) indicating that this sequence acts as a dominant ER-export motif and explaining why C-terminal fusions of E are retained in the ER.

**Figure 2.**
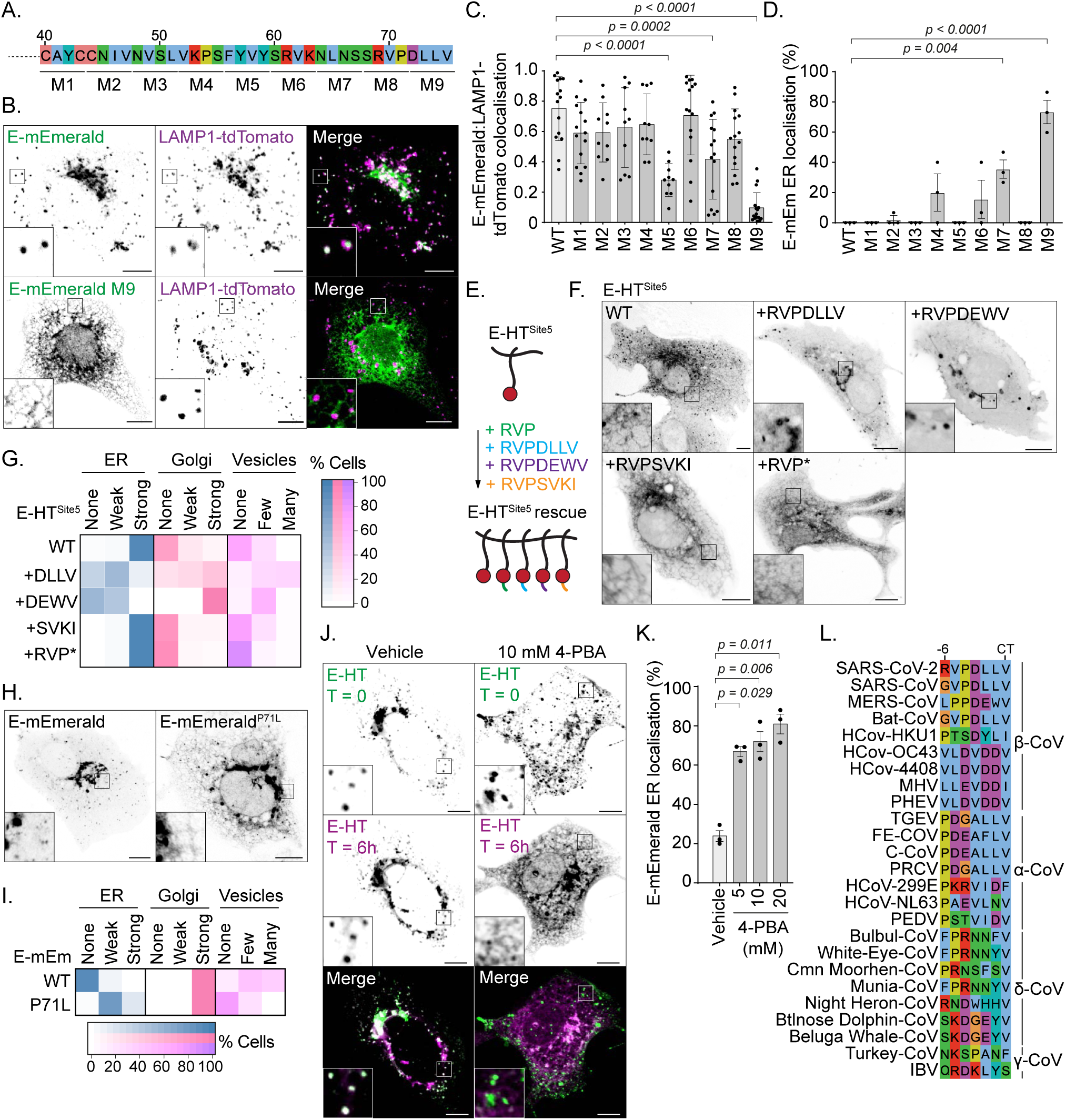
Alanine-scanning mutagenesis reveals C-terminal sequences necessary for ER-to-Golgi trafficking of E. (**A**) Schematic of alanine-scanning mutagenesis of SARS-CoV-2 E’s C-terminus. In mutants M1 – M9, this indicated amino acids were exchanged for Alanine. (**B**) Representative images of VeroE6 cells expressing the indicated versions of E-mEmerald and LAMP1-tdTomato. (**C, D**) Quantification of subcellular distribution of alanine-scanning mutants of E-mEmerald. Overlap of E-mEmerald with LAMP1-tdTomato was assessed by Manders’ Correlation Coefficient (M2) from 15 imaged cells with data points displaying M2 coefficients from all non-Golgi Envelope-positive regions of each cell, with mean ± S.D. displayed (C). ER localisation (D) was scored visually from 15 imaged cells in 3 independent experiments with mean ± S.E displayed. For C and D, statistical significance was determined by one-way ANOVA comparing each mutant against WT, with Dunnett’s correction for multiple testing. (**E, F**) Cartoon depiction of rescue assay (E) and representative images (F) of VeroE6 cells transfected with plasmids encoding the indicated JF646-illuminated C-terminal E-HT^Site5^ fusions followed by additions of the indicated chimeric terminal peptides from SARS-CoV-2 (RVPDLLV), MERS-CoV (RVP*DEWV*) or a Class-II PDZ ligand (RVP*SKVI*). Chimeric sequences italicised. (**G**) Quilt displaying localisation from 50 imaged cells for each condition in F. (**H, I**) Representative images of VeroE6 cells transfected with E-mEmerald or E-mEmerald^P71L^ with subcellular localization quantified (I) from 50 imaged cells. (**J**) Representative images of VeroE6 cells transfected with E-HT and illuminated with Oregon Green HaloTag ligand (green), released into dye-free media in the presence or absence of 4-PBA for 6 hours and then re-stained with TMR HaloTag ligand to illuminate newly-synthesised E-HT (magenta). (**K**) Quantification of ER localisation of newly synthesised TMR labelled E-HT from 50 imaged cells in 3 independent experiments imaged in (J). Mean ± S.E displayed, statistical significance determined by one-way ANOVA comparing each treatment to vehicle control, with Dunnett’s correction for multiple testing. (**L**) Sequence alignment of the extreme C-terminus of α, β, γ, and δ coronaviral E proteins. In microscopy panels, scale bars are 10 μm.

The C-terminal 4 amino acids of SARS-CoV E have been described to encode a PDZ-ligand^19^. This sequence is conserved in SARS-CoV-2 E and we wondered if engagement with a PDZ-domain containing partner licensed ER-export of E. However, we found that sequences from a variety of different classes of PDZ ligands could substitute for the DLLV sequence and drive ER-export, although none were as effective as chimeric C-termini from MHV (strain S) or MERS-CoV (Fig. S3A and S3B). The variety of ER-export competent PDZ-ligands from different classes argues against a specific PDZ-domain containing protein being required for ER export. Consistent with models of COP-II-dependent ER-export, the C-terminal valine of E provided most of the export activity, as E-mEmerald^ΔV^ was largely retained in the ER and exchanging the terminal DLLV for AAAV (E-mEmerald^DLLV-AAAV^) restored ER export (Fig S3C and S3D). However, a small pool of E-mEmerald^ΔV^ still reached the Golgi, suggesting that the context of this hydrophobic valine is important for ER-export. The beta-variant (B.1.351) of SARS-CoV-2 encodes E^P71L^ and we wondered whether this mutation influenced the efficiency of ER-export. Whilst E-mEmerald^P71L^ displayed steady state localisation to the Golgi, a fraction was retained in the ER and its ability to reach post-Golgi structures was limited (Fig. 2H and Fig. 2I), suggesting that impaired ER-export may be a feature of some previously circulating variants of SARS-CoV-2. C-terminal hydrophobic ER-export signals in secretory cargo proteins are typically recognised by the B-site of SEC24 isoforms^20^ for incorporation into the COP-II coat. Using a pulse-chase assay with sequentially applied HaloTag ligands, we found that 4-PBA, a small molecule that occludes the SEC24 B-site^21^, suppressed ER-export of newly synthesised E-HT (Fig. 2J and 2K). C-terminal hydrophobic residues are a conserved feature of E proteins (Fig. 2L), suggesting that SEC24-dependent ER-export may be utilised across *coronaviradae* to access the Golgi for viral assembly.

### Identification of host-factors interacting with E by proximity biotinylation

We inserted a hemagglutinin (HA)-tagged TurboID^22^ into the internal tagging sites in E and confirmed that this did not disrupt E’s localisation (Fig. S4A). After confirming that versions of E-HA/TurboID co-localised with E-mEmerald in 293T cells (Fig S4B), we used proximity biotinylation, mass spectrometry (MS) and label-free quantification (LFQ) to determine the proximal proteome of E in these cells (Fig. S4C-S4E). We found that many ERGIC and Golgi proteins, and components of both anterograde (SEC24B) and retrograde (RER1, COP-epsilon) transport machineries were significantly enriched by E-HA/TurboID, relative to a cytosolic control (Fig. S4E). We confirmed physical interactions with RER1, GORASP2 and PALS-1, a previously identified SARS-CoV E and SARS-CoV-2 E interacting partner^23,24^ (Fig. S4F). We next compared proximal proteomes from ER-export proficient (E-HA/TurboID^Site3^, E-HA/TurboID^Site4^, E-HA/TurboID^Site3ΔDLLV+DEWV^) and ER-export defective (E-HA/TurboID^Site3ΔDLLV^, E-HA/TurboID^Site4ΔDLLV^ and E-HA/TurboID^Site3ΔDLLV+SVKI^) versions of E. Reported proximal proteomes from these differentially localised versions of E clustered well by Principal Component Analysis and hierarchical clustering (Fig. S5A – S5C). We recovered peptides from numerous PDZ-domain containing proteins with WT but not ΔDLLV versions of E-HA/TurboID (Fig. S5D and S5E), confirming that this sequence can act as a PDZ-ligand. We observed strong enrichment of Golgi and ERGIC proteins for ER-export competent versions of E-HA/TurboID, and strong enrichment of ER-proteins for versions of E lacking the ability to escape the ER (Fig. S5E). Consistent with our identification of COP-II-dependent ER-export (Fig. 2), we recovered the COP-II components SEC24A, SEC24B and SEC31A with ER-export competent versions of E (Fig S5E). Lastly, in agreement with our imaging approaches documenting the localisation of E to lysosomes, we detected significant enrichment of endosomal and lysosomal proteins in our ER-export-competent versions of E (Fig. S5E). We identify here an extensive set of interaction partners for SARS-CoV-2 E across biosynthetic and endocytic pathways.

### Identification of machinery necessary for Golgi-to-lysosome trafficking of SARS-CoV-2 Envelope

We next questioned how E was delivered to lysosomes. Whilst there exist direct trafficking pathways between Golgi and endosomes, some lysosomal proteins are first delivered to the cell surface and then internalised via endocytic routes to allow lysosomal localisation. Alternatively, the heterotetrameric clathrin adaptor complex AP-1 can select cargo for *trans*-Golgi Network (TGN)-to-endosome transport, where it works in-concert with the Golgi-localised Gamma-ear-containing Adaptor-1 (GGA1) and AP1AR/Gadkin, a kinesin adaptor responsible for the anterograde movement of AP-1 carriers^25–27^. An AP-3 dependent pathway is also thought to deliver cargo directly from Golgi to lysosomes, although this is less well characterised in mammalian cells^28^. Expression of a dominant negative form of the endocytic GTPase, Dynamin^29^, robustly blocked transferrin internalisation but had no impact on the intracellular distribution of E-mEmerald (Fig. S6A and S6B), suggesting that E is not trafficked to lysosomes via the plasma membrane. To investigate host-cell factors responsible for Golgi-export of E, we selected 12 membrane trafficking genes identified as high-confidence hits from our proximal proteome (Fig S6D) and used CRISPR-Cas9 to delete them in VeroE6 cells. We verified homozygous deletion for each target by next-generation sequencing or western blotting (Fig. 3A, Fig. S7A and Fig. S7B) and compared localisation of E-mEmerald in these lines (Fig. S7C and Fig. S7D). In ARFRP1^-/-^ cells, we found that whilst E-mEmerald was able to exit the ER it was retained in TGN46-positive tubules emanating from the TGN and did not reach lysosomes (Fig. 3B, Fig. S7C and Fig. S7D). Importantly, transmembrane proteins such as LAMP1 were correctly localised in ARFRP1^-/-^ cells (Fig. 3B), suggesting that these cells do not exhibit a global block in Golgi export. ARFRP1 is an ARF1-related GTPase. Endogenous ARFRP1 colocalised with GFP-TGN46 and ARFRP1-positive membranes were juxtaposed against membranes positive for the *cis*- and *medial*-cisternae localising golgin, Giantin (Fig 3C), indicating that ARFRP1 is a TGN-resident enzyme. We next re-expressed versions of ARFRP1 in ARFRP1^-/-^ VeroE6 cells to test requirements for its enzymatic activity in the Golgi export of E-mEmerald. Re-expression of ARFRP1 or its catalytically active mutant, ARFRP1^Q79L^, in ARFRP1^-/-^ cells could restore Golgi export of E-mEmerald and suppress its retention in TGN-derived tubules. Re-expression of a dominant-negative ARFRP1^T31N^ or ARFRP1^Y89D^, a version of ARFRP1 containing a mutation in its hydrophobic effector patch^30^ (equivalent to ARF1^Y81D^, Fig. S7E), could not (Fig. 3D). Interestingly, ARFRP1^Y89D^ and ARFRP1^T31N^ still localised to the Golgi, but were not themselves incorporated into the E-mEmerald-containing TGN-derived tubules (Fig 3D), suggesting that ARFRP1 coordinates a machinery allowing carrier formation for Golgi-to-endosome trafficking of E. We interrogated our E-HA/TurboID proximal interactome and noted enrichment of members of the AP-1 clathrin adaptor complex, AP1AR/Gadkin and GGA1 (Fig. S8A). AP-1 has been identified as necessary for SARS-CoV-2 replication in several genome wide CRISPR screens^31–33^, but the mechanistic basis for its contribution to the SARS-CoV-2 lifecycle remains unexplored. AP-1 could be detected as puncta on ARFRP1-positive E-mEmerald-positive TGN membranes (Fig. 3E and Fig. S8B). Loss of AP-1 via siRNA-mediated depletion of AP1M1 phenocopied ARFRP1-deletion, with E-mEmerald retained in TGN-derived tubular structures, suggesting that AP-1 and ARFRP1 operate in the same pathway to allow Golgi export of E (S8C-S8E). Interestingly, depletion of AP1AR/Gadkin in ARFRP1^-/-^ cells suppressed formation of these tubules (Fig. S8C-S8E), suggesting that they are generated via coupling to anterograde microtubule motors. Finally, GGA1-depletion mimicked the loss of AP1M1 and similarly led to the retention of E-mEmerald in Golgi-derived tubular structures (Fig. S8C-S8E), suggesting it operates alongside ARFRP1 and AP-1 in the export of E-mEmerald from the Golgi. In the case of depletion of AP1M1 or GGA1 in ARFRP1^-/-^ cells, we observed no additive phenotypes in tubule formation (Fig. S8E), suggesting that these proteins operate in the same pathway. Given the similarities in E-mEmerald phenotypes produced upon the inactivation of ARFRP1 and AP-1, we examined AP-1 localisation in ARFRP1^-/-^ cells and discovered that it was delocalised from the TGN (Fig. 3F and Fig 3G). AP-1 localisation to the TGN could be restored in ARFRP1^-/-^ cells by re-expression of ARFRP1 or ARFRP1^Q79L^, but not by re-expression of ARFRP1^T31N^ or ARFRP1^Y89D^ (Fig. 3H and 3I). Finally, we found that overexpression of ARFRP1^T31N^ or ARFRP1^Y89D^ could delocalise endogenous AP-1 from the TGN, suggesting that these mutants act as dominant negative inhibitors of AP-1 at this organelle (Fig. S8F and S8G). These data identify ARFRP1 as a TGN-localised GTPase whose activity is necessary for localising AP-1 to this organelle, and reveal that an ARFRP1/AP-1 dependent pathway allows TGN export of E and delivery to lysosomes.

**Figure. 3.**
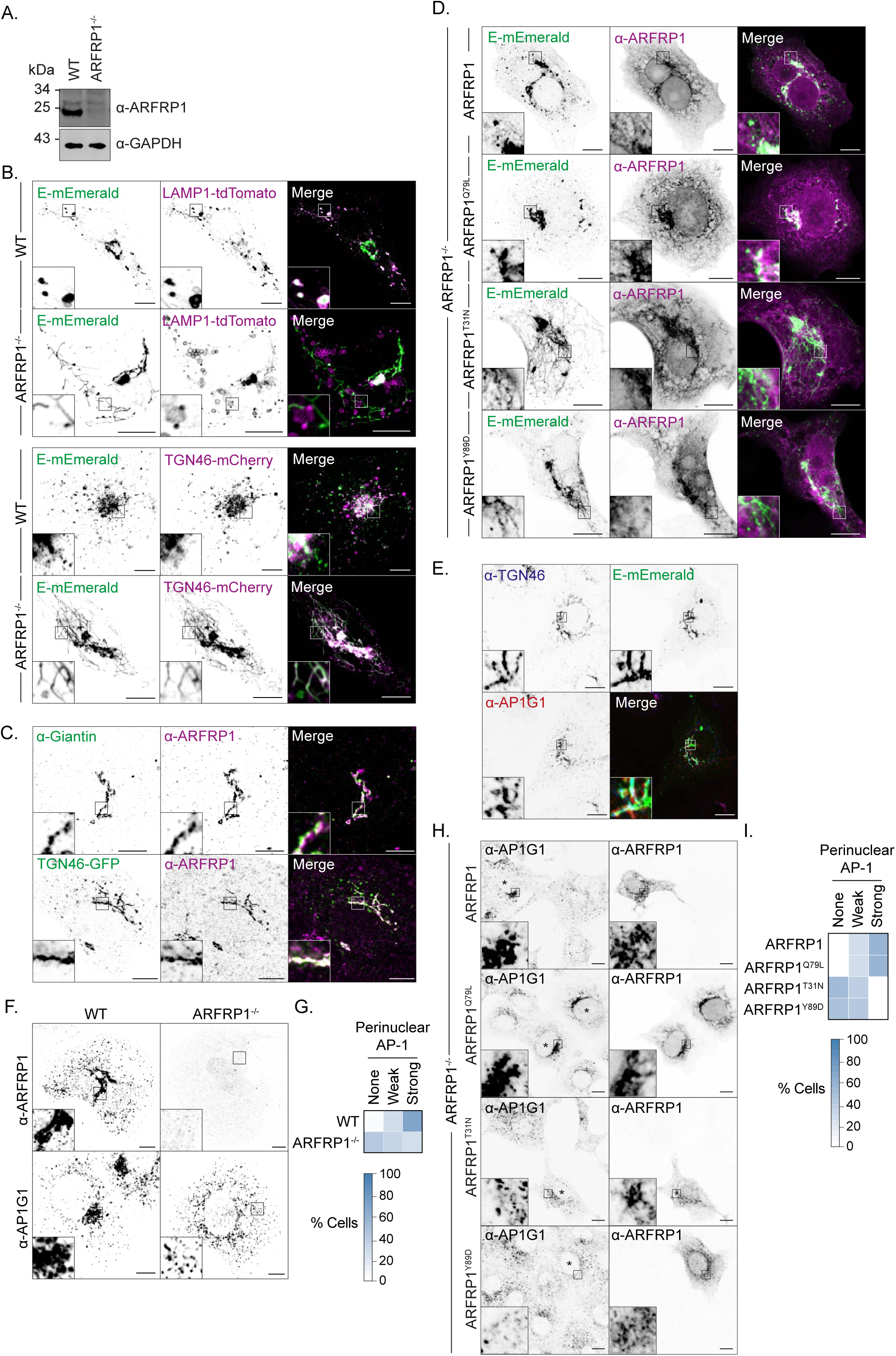
An ARFRP1/AP-1 axis controls Golgi-to-lysosome delivery of SARS-CoV-2 Envelope. (**A**) Resolved lysates from WT or ARFRP1^-/-^ VeroE6 cells were examined by western blotting with antibodies raised against ARFRP1 or GAPDH. (**B)** Representative images of WT or ARFRP1^-/-^ VeroE6 cells transfected with plasmids encoding E-mEmerald and either LAMP1-tdTomato or TGN46-mCherry. Images representative of 15 imaged cells in each case. (**C**) VeroE6 cells or VeroE6 cells transfected with a plasmid encoding TGN46-EGFP were fixed and stained with antisera raised against Giantin or ARFRP1. Images representative of 15 imaged cells in each case. (**D**) Representative images of ARFRP1^-/-^ VeroE6 cells transfected with plasmids encoding E-mEmerald and either ARFRP1, ARFRP1^Q79L^, ARFRP1^T31N^ or ARFRP1^Y89D^. Cells were stained with antibodies raised against ARFRP1 to detect transfected cells. Images representative of 15 imaged cells in each case. (**E**) VeroE6 cells were transfected with a plasmid encoding E-mEmerald and stained against endogenous AP1G1 and TGN46. (**F, G**) WT or ARFRP1^-/-^ VeroE6 cells were fixed and stained with antisera raised against ARFRP1 or AP1G1 and perinuclear localisation of AP1G1 was scored in the accompanying quilt from 50 imaged cells (G). (**H, I**) Plasmids encoding the indicated ARFRP1 proteins were transfected into ARFRP1^-/-^ VeroE6 cells. Cells were fixed and stained with antisera raised against ARFRP1 or AP1G1 and the perinuclear localisation of AP1G1 was scored in the accompanying quilt from over 50 imaged cells per condition across 3 independent experiments (I). Transfected cells indicated by asterisks. In microscopy panels, scale bars are 10 μm.

### SARS-CoV-2 Envelope binds AP-1 via sequences required for Golgi-export

We next returned to our alanine-scanning mutagenesis to explore viral sequences necessary for Golgi export of E. We noted that E-mEmerald^M5^ was exported well from the ER but was not delivered from the Golgi to lysosomes, and was instead retained in tubules emanating from this organelle (Fig. 4A and Fig. 4B, Fig. S2 and Fig. S6C). Adaptins recognise cargos by binding Short Linear Interaction Motifs (SLIMs) presented in the cytosolic region of transmembrane cargos. SLIMs including YxxΦ and FxxFxxxR are recognised by hydrophobic pockets in Mu-2 and Beta-2 adaptins, respectively^34–36^. These hydrophobic pockets are well conserved in Mu-1 and Beta-1 adaptins, and we noted similarities between the sequences surrounding the residues mutated in E-mEmerald^M5^ that were necessary for Golgi export, and these SLIMs (Fig 4A). We mutated either Y59A and F56A/Y59A/K63A (E-mEmerald^Y59A^ and E-mEmerald^FYK-AAA^) to disrupt these putative AP-1 interactions and examined lysosomal delivery of E-mEmerald. Whilst E-mEmerald^Y59A^ was delivered normally to lysosomes, we found that E-mEmerald^FYK-AAA^ was retained in TGN-derived tubular carriers, mimicking the effects of E-mEmerald^M5^, or the effects of inactivating ARFRP1 or AP-1 (Fig. 4B-4D). The distribution of hydrophobic and basic residues in this region (residues 56-63) is well conserved amongst β-coronaviruses but is absent from α-coronaviruses (Fig 4E). We deleted this sequence from E-mEmerald and exchanged it with equivalent sequences from either β-coronaviral (MERS-CoV and OC43), or α-coronaviral (hCov299 or TGEV) E proteins (Fig. 4F). Confirming requirements for this region in Golgi export, E-mEmerald^Δ56–63^ was retained in Golgi-derived tubules (Fig. 4G and 4H). Delivery to lysosomes was rescued by insertion of equivalent sequences from β-coronaviral, but not α-coronaviral, E proteins (Fig 4G and 4H). We used GFP-Trap co-precipitation assays to test interaction with AP-1. We found that E-mEmerald could bind HA-tagged and endogenous AP1B1, (Fig. 4I and 4J), that deletion of residues 56-63 reduced the interaction with HA-AP1B1, and that this binding could be rescued using chimaeric sequences from β-coronaviral, but not α-coronaviral, E proteins (Figure 4J and 4K). These data identify an ARFRP1- and AP-1-dependent membrane trafficking pathway that exports E from the Golgi to lysosomes, identify viral sequences that bind AP-1 and demonstrate conservation of these properties amongst β-coronaviral, but not α-coronaviral, E proteins.

**Figure 4.**
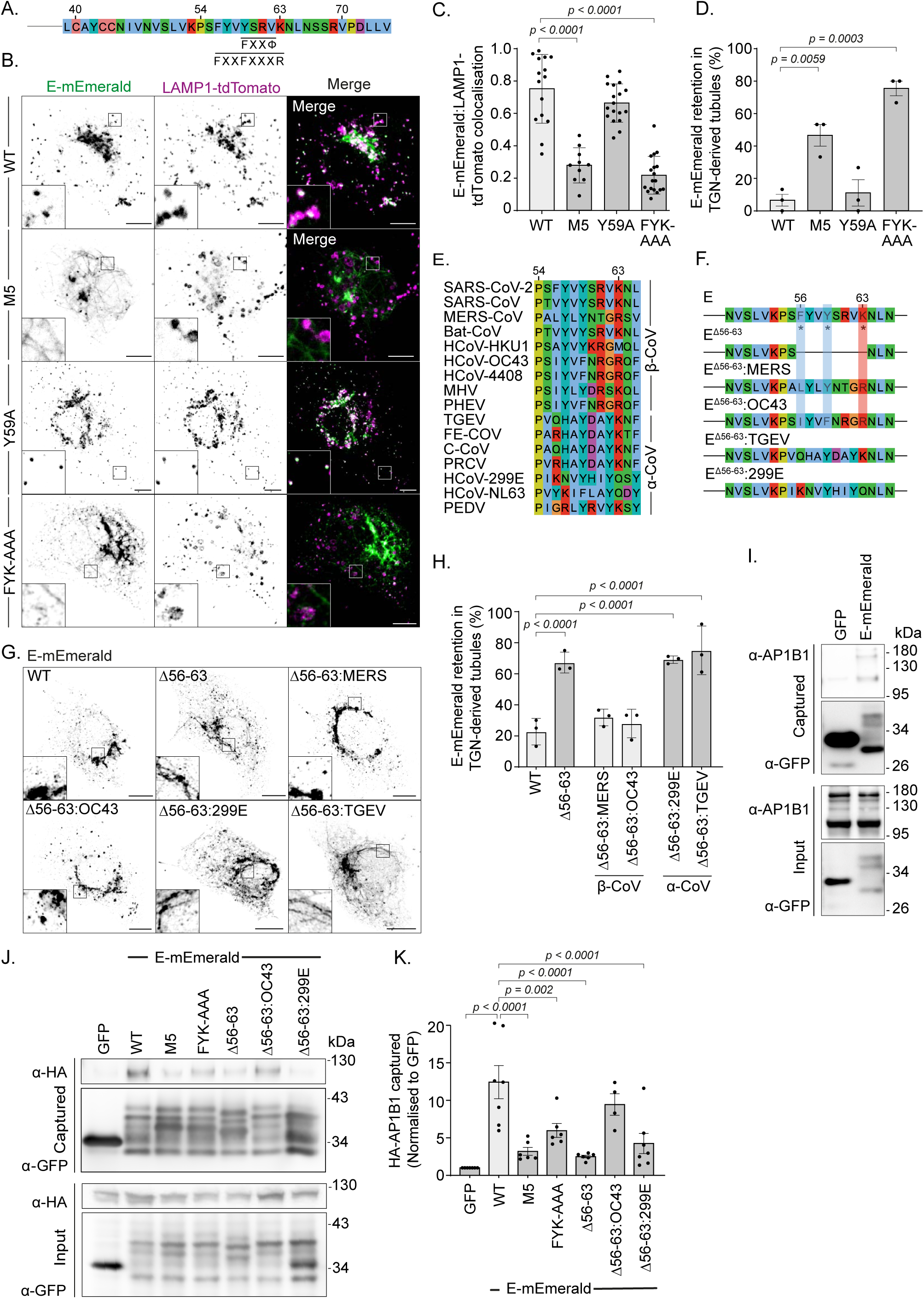
SARS-CoV-2 Envelope binds the AP-1 adaptor protein complex. (**A**) Schematic depiction of the SARS-CoV-2 E’s cytosolic C-terminus with putative AP-binding SLIMs highlighted. (**B-D**) Representative images of VeroE6 cells transfected with plasmids encoding the indicated E-mEmerald plasmids and LAMP1-tdTomato (B) with quantification of the overlap of E-mEmerald with LAMP1-tdTomato as assessed by Manders’ Correlation Coefficient (M2) from 15 imaged cells (C). Data points in C represent M2 coefficients from all non-Golgi Envelope-positive regions of each cell, with mean ± S.D. displayed. Tubular E-mEmerald localisation (D) was scored visually from 15 imaged cells in 3 independent experiments with mean ± S.E displayed. For C and D, statistical significance was determined by one-way ANOVA comparing each mutant against WT, with Dunnett’s correction for multiple testing. (**E**) Sequence alignment of the cytosolic region of E responsible for Golgi-to-lysosome trafficking and the equivalent region from β- and α-coronaviruses. (**F**) Schematic of E chimaeras in which the sequence between 56-63 was deleted and/or replaced with the equivalent sequences from β-coronaviruses (MERS, OC43) or α-coronaviruses (TGEV, 299E). (**G, H**) Representative images of VeroE6 cells transfected with plasmids encoding the indicated E-mEmerald (G) and quantification of cells displaying a retention of E-mEmerald in Golgi-derived tubular structures (H) from 15 imaged cells and 3 independent experiments. Mean ± S.E is displayed, with significance determined by one-way ANOVA with Šidák’s correction for multiple testing. (**I**) Cell lysates and GFP-Trap immunoprecipitations from 293T cells transfected with plasmids encoding GFP or E-mEmerald were resolved by SDS-PAGE and examined by western blotting with antisera raised against GFP or AP1B1. (**J**) Cell lysates and GFP-Trap immunoprecipitations of 293T cells co-transfected with plasmids encoding GFP or the indicated E-mEmerald constructs and HA-AP1B1 were resolved by SDS-PAGE and examined by western blotting with antisera raised against HA or GFP. (**K**) Quantification of data from J from N = 4 to 7 independent experiments as indicted by individual data points. In microscopy panels, scale bars are 10 μm.

### The role for endolysosomal trafficking of E in the SARS-CoV-2 replication cycle

*Trans*-complementation assays have proved powerful for understanding viral elements necessary for replication in a variety of systems^37,38^. To understand how both ER-export and ARFRP1/AP-1-dependent delivery of E from Golgi to lysosomes contributes to the SARS-CoV-2 replication cycle, we developed an RNA-interference strategy allowing targeting of the sub-genomic SARS-CoV-2 RNA responsible for producing E. Like other *Nidovirales*, SARS-CoV-2 employs discontinuous transcription during negative-strand RNA synthesis to allow template switching between transcription-regulating sequences (TRS) in the leader sequence of *ORF1A/B* (TRS-L) and identical sequences (TRS-B) immediately upstream of ORFs in the 3’-end of the genome^39–41^ (Fig. 5A). This allows production of the sub-genomic RNAs (sgRNA) encoding S, E, M, N and several non-structural proteins. Using firefly and renilla luciferase reporters, we designed siRNAs targeting the TRS/E junction (Fig. S9A and Fig. S9B) to deplete sgRNA encoding E. We identified sequences that targeted E sgRNA but spared both genomic SARS-CoV-2 RNA and N sgRNA (Fig. S9C and S9D) and used qRT-PCR to confirm these oligos were able to target E sg-mRNA, but not N sg-mRNA, in the context of a viral infection (Fig. S9E). We next used the *Sleeping-Beauty* retrotransposition system (Fig. S9F) to integrate a cassette encoding a constitutively expressed tdTomato and a doxycycline-inducible codon-optimised version of E into VeroE6 cells to allow *trans*-complementation of E. We verified dox-inducible expression of E-mEmerald in equivalently transposed VeroE6 cells sorted on tdTomato (Fig S9G), generated equivalently sorted versions of VeroE6 cells expressing codon-optimised doxycycline-inducible versions of E, E^N15A/V25F^, E^M5^, E^FYK-AAA^, or E^ΔDLLV^ and transfected them with E-targeting siRNA. After 20 hours, we infected these cells with SARS-CoV-2 (hCoVL19/England/02/2020) and assessed the titer of virus produced via *trans*-complementation by using plaque assay. As expected, depletion of E attenuated, although did not eliminate, the amount of infectious SARS-CoV-2 produced (Fig S9H-J). This could be rescued robustly by *trans*-complementation with wild-type E but not by versions of E that were retained in the ER (E^ΔDLLV^) or could not be exported from the Golgi and delivered to lysosomes (E^M5^, E^FYK-AAA^) in the producer cell (Fig. 5B-5D). Importantly, *trans*-complementation with a version of E containing mutations that abrogate its viroporin activity (E^N15A/V25F^)^42^ did not rescue viral titers in this system (Fig. 5B-5D). These data provide functional evidence that intracellular trafficking of E from both ER-to-Golgi and from Golgi-to-lysosomes in host cells supports SARS-CoV-2 replication. Consistent with other systems in which recombinant β-coronaviruses lacking E produce smaller and irregularly shaped plaques^13^, *trans*-complemented versions of SARS-CoV-2 bearing versions of E with disrupted viroporin activity, or that were unable to reach lysosomes, produced smaller plaques (Fig 5D). Finally, to distinguish entry and release effects, we turned to a Virus Like Particle (VLP) system to examine roles for E in particle assembly and release. Using a 4-component (E, S, M, N) SARS-CoV-2 VLP system, we found that particle release was impaired when we used versions of E that were either defective in their viroporin activity (E^N15A/V25F^) or that could not be trafficked from Golgi to lysosomes (E^M5^) (Fig 5E-5G). We also found that particle production was permissible using versions of E that were restricted to the ER (E^ΔDLLV^), but in this case, packaging of N was impaired, suggesting that the site of assembly allows proper biogenesis of SARS-CoV-2 particles (Fig. 5E-5G). These data suggest that trafficking of E as a functional viroporin to lysosomes contributes to late stages of the SARS-CoV-2 replication cycle.

**Figure. 5.**
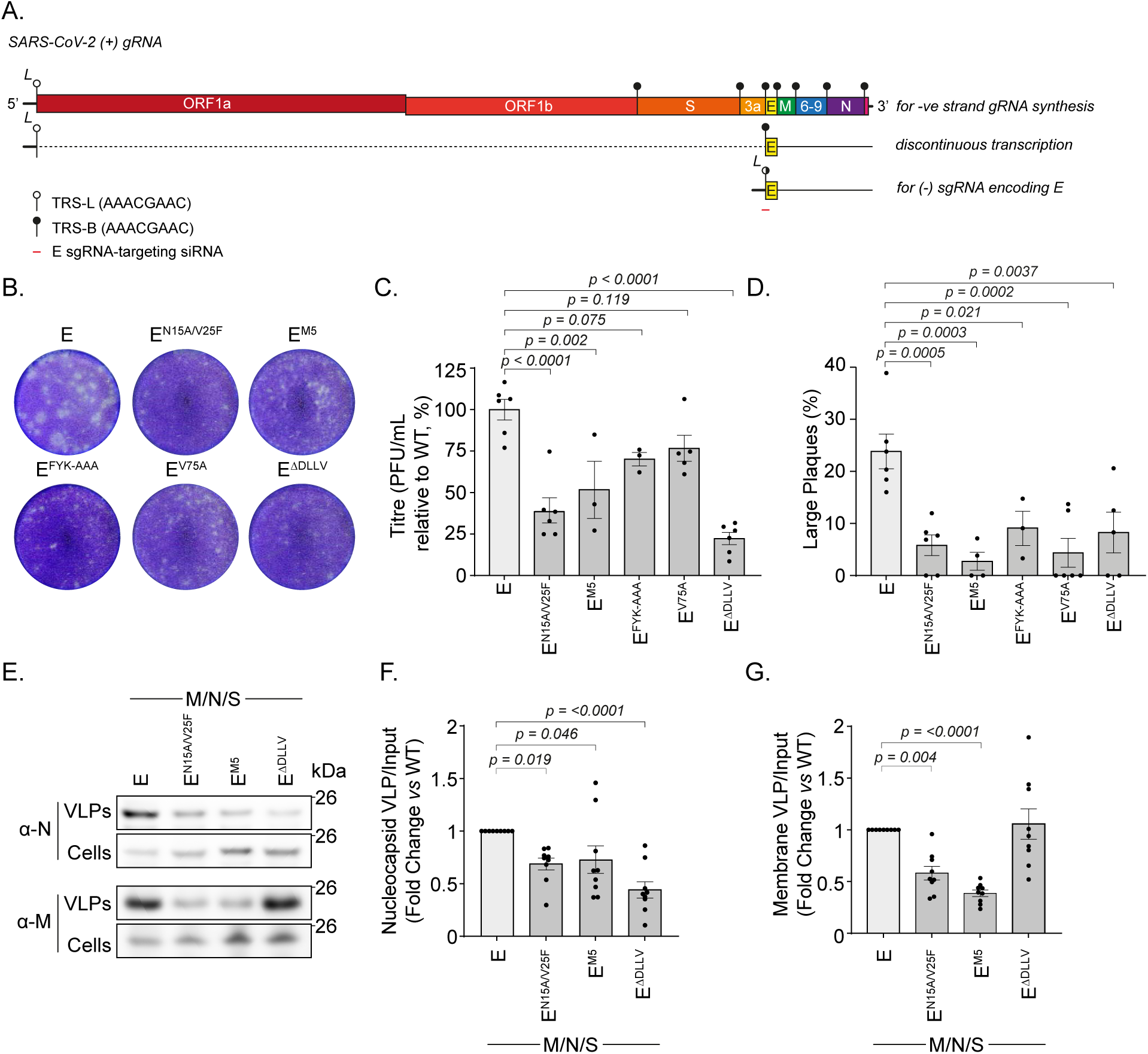
SARS-CoV-2 Envelope trafficking mutants disrupt viral egress. (**A**) Schematic of transcription regulatory sequences (TRS) in the SARS-CoV-2 genome, the discontinuous transcription of sgmRNAs, and the design location of E-sgmRNA siRNAs. (**B**) Supernatants from VeroE6 cells containing the indicated dox-inducible codon-optimised E constructs, that had been transfected with E-sgmRNA-targeting siRNA, treated with doxycycline and infected with SARS-CoV-2 (hCoVL19/England/02/2020) were used to infect fresh VeroE6 cells, and plaques were allowed to develop for 3-days before being fixed and stained using 0.2 % toluidine blue. Images show representative plaque formation. (**C**) Quantification of plaque formation represented as titre (PFU/ml) using the ViralPlaque FIJI macro. Mean ± S.E. presented with significance calculated by a 1-way ANOVA with Dunnet’s correction applied for multiple testing. (**D**) Quantification of the percentage of large plaques *vs* total plaques from plaque assays. Large plaques were defined by having an area greater than 0.82 mm^2^ and measured using the ViralPlaque macro in FIJI. Mean ± S.E. presented, with significance calculated by a 1-way ANOVA with Dunnet’s correction applied for multiple testing. N = 3 to 6 independent experiments, as indicated by data points. (**E**) Resolved cell lysates and VLP fractions from 293T cells transfected with the indicated codon-optimised versions of M, N, S and E were examined by SDS-PAGE and immunoblotted using antisera raised against SARS-CoV M and SARS-CoV-2 N. (**F, G**) Quantification of N or M present in VLPs generated using either WT or mutant versions of E normalised against N or M present in cell lysates. Data plotted as fold change relative to WT. Mean ± S.E. presented from N = 9, with significance calculated by a 1-way ANOVA with Dunnet’s correction applied for multiple testing.

## DISCUSSION

We have demonstrated that the small Envelope protein from SARS-CoV-2 encodes sequence specific information that allows it to navigate the host’s endomembrane network, is routed to lysosomes and acts as a viroporin to neutralize the pH in these organelles. We found that E encodes an ER-export sequence, mediated primarily by a C-terminal hydrophobic residue and engagement with COP-II vis SEC24. C-terminal hydrophobic residues are conserved across α-, β- and γ-coronaviruses, pointing to a conserved mechanism of ER-export for E proteins across *coronaviridae*. Previously published C-terminally tagged versions of E localise inappropriately to the ER^43^, and we suggest here that this is due to occlusion of this dominant ER-export sequence.

Secondly, our internally tagged versions of E allowed us to report that whilst the majority of E localises to the Golgi, a pool of E is delivered from here to lysosomes. We identified sequence motifs within the cytosolic C-terminus of E that allow Golgi-to-endosome trafficking and we exposed a role for ARFRP1 in coordinating an AP1- and AP1AR/Gadkin-dependent pathway that allows trafficking of E from Golgi to lysosomes. Of note, AP-1 and AP1AR/Gadkin have been implicated previously in the release of cargo by secretory lysosomes^25^. ARFRP1 is needed for both the recruitment of golgins and GARP to the TGN^44^ and controlling export of the planar cell polarity protein, Vangl2^45^. We show here that ARFRP1 is a TGN-localised GTPase whose GTPase activity is necessary for both AP-1 recruitment to the Golgi and for Golgi-to-endosome trafficking of E. AP-1 plays a complex role in bi-directional traffic between the Golgi and endosomes, acting alone in the retrograde pathway from endosomes-to-Golgi, and in-concert with GGAs and AP1AR/Gadkin in the anterograde pathway from Golgi-to-endosomes^28^. Consistent with role AP-1 in the anterograde movement of E, when this pathway was inactivated, we observed a failure of Golgi export, rather than a redistribution of E to endosomes. We show that the Golgi-retention phenotype of E-mEmerald^M5^ is attributable to the loss of AP-1 binding, and we show that sequences required for AP-1-binding and for Golgi-to-endosome trafficking are conserved within β-coronaviruses, but not α-coronaviruses. These data suggest that β-coronaviral E proteins have evolved to exploit an ARFRP1/AP-1-dependent trafficking pathway for transport between Golgi and endosomes, and provide context to the identification of AP-1 in genome wide screens for host factors regulating SARS-CoV-2 replication.

Consistent with the work of others, and findings in SARS-CoV-2 infected cells^16,17^, we found that E was able to neutralise lysosomal pH. Whilst the ORF3a proteins of SARS-CoV or SARS-CoV-2 have been proposed as ion channels^46^, recent cryo-EM and electrophysiological evidence suggests that these proteins do not act as viroporins^47^, and that they impose their effect on lysosomal biology through interaction with the HOPS complex^47,48^. Given the potential for viral egress through deacidified secretory lysosomes and the finding that lysosomal pH is neutralized in SARS-CoV-2 infected cells^15,16^, we suggest that trafficking of E to lysosomes contributes significantly to the neutralization of pH in this organelle.

What role does lysosomal pH neutralization play in the SARS-CoV-2 lifecycle? In SARS-CoV systems, E’s channel activity was necessary for viral pathogenesis, with recombinant viruses bearing channel mutations acquiring compensatory mutations to restore ion flux^42^. The omicron variant of SARS-CoV-2 encodes a version of E with a point mutation (E^T9I^) in a polar channel-lining residue is less able to neutralise lysosomal pH which contributes to a reduced viral load in SARS-CoV-2 infected cells^17^. That the combined channel and oligomerization mutant of E (E^N15A/V25F^) was poorly able to support SARS-CoV-2 replication when supplied in-*trans* suggests that channel activity is necessary for a productive infection. We reasoned that neutralization of lysosomal pH would either limit exposure of internalized virus to this proteolytic compartment, or could protect virions in secretory lysosomes from this degradative environment. Our VLP production assays allowed examination of egress effects, and our findings that E^N15A/V25F^ or E^M5^ reduced VLP release suggests that these deacidified lysosomes are important for preserving particles during egress. We also note that our VLP assays also showed that packaging of N was most efficient when E contained an intact C-terminus. Whilst in SARS-CoV, N has been proposed to interact with the extreme C-terminus of E^49^, the reduced incorporation of N into VLPs containing E^ΔDLLV^ may also be a consequence of restricted the localisation of E to the assembly site.

Finally, our proximal interactomes provide a powerful resource for understanding host factors that may regulate E’s biology. Notably, we recovered many new PDZ-domain containing proteins that were biotinylated in a manner requiring E’s extreme C-terminus, many of which have been subsequently validated, including PALS1^23^ and TJP1^50^ and which likely contribute to epithelial barrier function. Our interactomes differ from those reported by affinity purification^1^, but do not suffer from high-level overexpression or placement of affinity or BioID tags that would disrupt the normal localisation of E^1,51–53^. Secondly, we present a *trans*-complementation assay allowing depletion and rescue of sub-genomic RNAs encoding SARS-CoV-2 structural proteins, allowing us to take reverse genetic approaches without needing to create genetically modified recombinant SARS-CoV-2 viruses. We anticipate that targeting the TRS elements for alternate sub-genomic RNAs will allow *trans*-complementation of these proteins in future.

In summary, our data have outlined trafficking pathways and routes taken by the E viroporin of SARS-CoV-2, linking viral sequences with cellular factors that govern movement between the ER, Golgi and lysosomes. We have uncovered new pathways responsible for the localization of AP-1 at the Golgi. We find specific effects of E on the neutralisation of lysosomal pH, which enables efficient particle release and SARS-CoV-2 replication. As well as facilitating viral egress, given the role of the lysosome as a terminal degradative organelle for a variety of cellular routes, we suspect that E’s expression will have wide ranging effects on the proteostatic capabilities of infected cells.

## Acknowledgements

J.G.C. is a Wellcome Trust Senior Research Fellow (206346/Z/17/Z & 224484/Z/21/Z) and is supported by the Francis Crick Institute which receives its core funding from Cancer Research UK (CC1002, CC2166), the UK Medical research Council (CC1002, CC2166), and the Wellcome Trust (CC1002, CC2166). We thank Rocco D’Antuono and the Crick Advanced Light Microscopy facility for access to equipment and FLIM analysis, and Dr Sharon Tooze (Crick) for access to her Zeiss LSM 880. This research was funded in whole, or in part, by the Wellcome Trust (206346/Z/17/Z, 224484/Z/21/Z, CC1002, CC2166). For the purpose of Open Access, the authors have applied a Creative Commons Attribution (CC BY) public copyright licence to any Author Accepted Manuscript version arising from this submission.

## Supplementary Figure Legends

**Figure S1.**
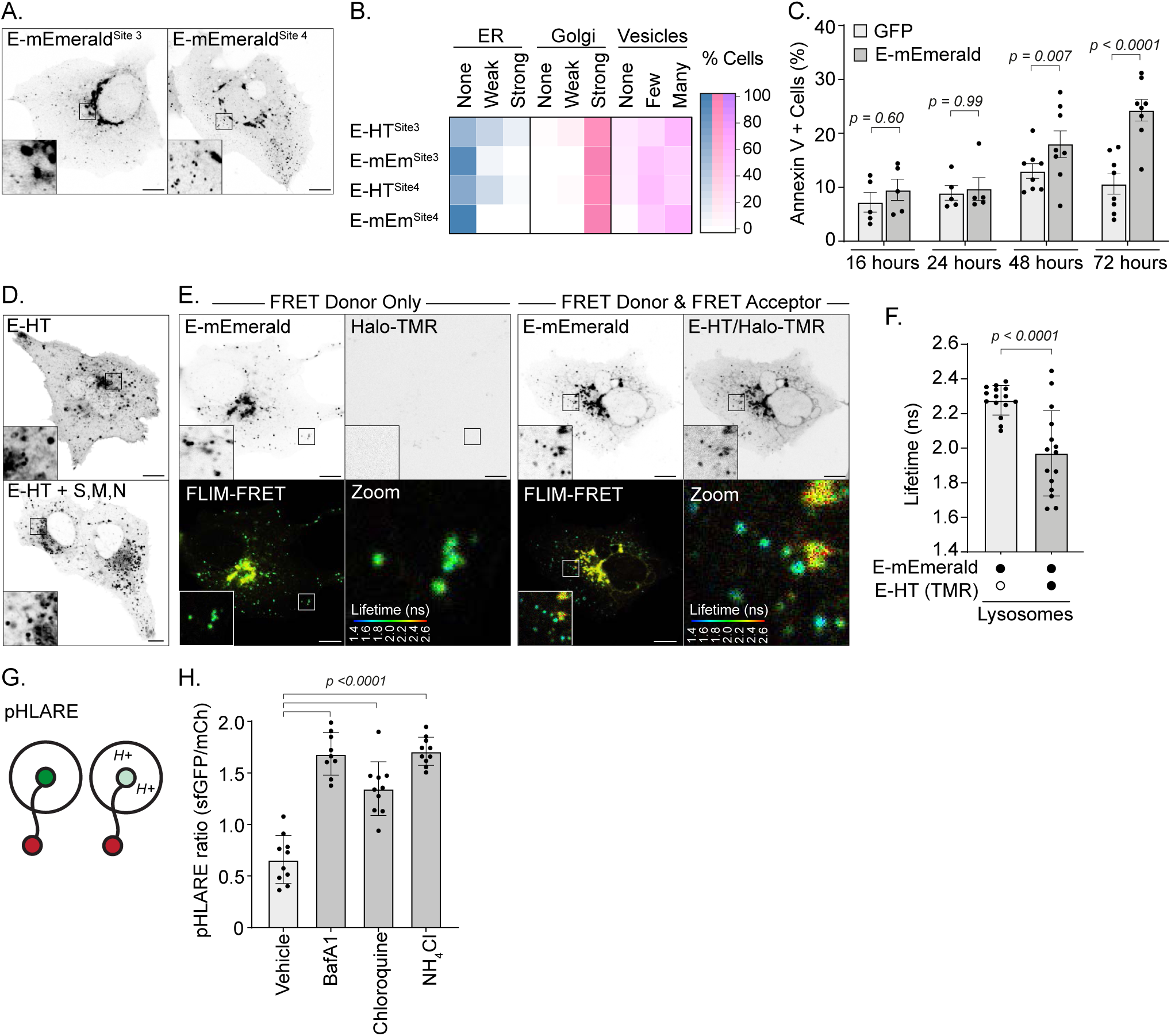
Validation of tagging strategy and ability of tagged versions of E to deacidify lysosomes. (**A**) Representative images of VeroE6 cells transfected with plasmids encoding the indicated mEmerald-fusions of E and imaged live. (**B**) Quilt of subcellular distribution of the indicated E-mEmerald or JF646-illuminated E-HT fusions from 50 imaged cells. (**C**) Flow cytometry-based analysis of Annexin V+ VeroE6 cells transfected with either GFP or E-mEmerald for the indicated times. At least 3000 GFP+ events captured per condition, N = 5 independent experiments for 16 hours and 24 hours, and N = 8 independent experiments for 48 hours and 72 hours. Mean ± S.E. displayed, with significance determined by 2-way ANOVA with Šidák’s correction for multiple testing. (**D**) Representative images of JF646-illuminated E-HT fusions transfected into VeroE6 cells either alone or with plasmids encoding SARS-CoV-2 Spike (S), Nucleocapsid (N) and Membrane (M) proteins. Images representative of 15 imaged cells. (**E**) Representative confocal and FLIM-FRET images of VeroE6 cells transfected with plasmids encoding E-mEmerald and E-HT. E-HT was illuminated with Tetramethyl Rhodamine (TMR). A rainbow LookUp-Table (LUT) was used to display mEmerald lifetime. (**F**) Quantification of the E-mEmerald lifetime in segmented lysosomal or Golgi classes. Mean ± S.D. displayed, with significance testing performed by 2-tailed T-Test from > 40 lysosomes in 15 imaged cells per condition. Average mEmerald lifetime in all JF646-positive lysosomes in each cell plotted as datapoints. (**G**) Schematic of pHLARE assay; pHLARE consists of a LAMP1 with N-terminal sfGFP and C-terminal mCherry tags with the pKa of sfGFP enabling quenching in an acidic environment. (**H**) VeroE6 cells were transfected with pHLARE and treated with the indicated compounds (Bafilomycin A1, 200 nM, 150 minutes; Chloroquine, 10 µM, 160 minutes; Ammonium Chloride, 10 mM, 180 minutes). Cells were fixed and imaged and the ratio of sfGFP to mCherry fluorescence in mCherry positive lysosomes was calculated. Mean ± S.D. presented from N = 10 cells per condition, significance calculated by 1-way ANOVA comparing treatments to vehicle control, with Dunnett’s correction applied for multiple testing. In microscopy panels, scale bars are 10 μm.

**Figure S2.**
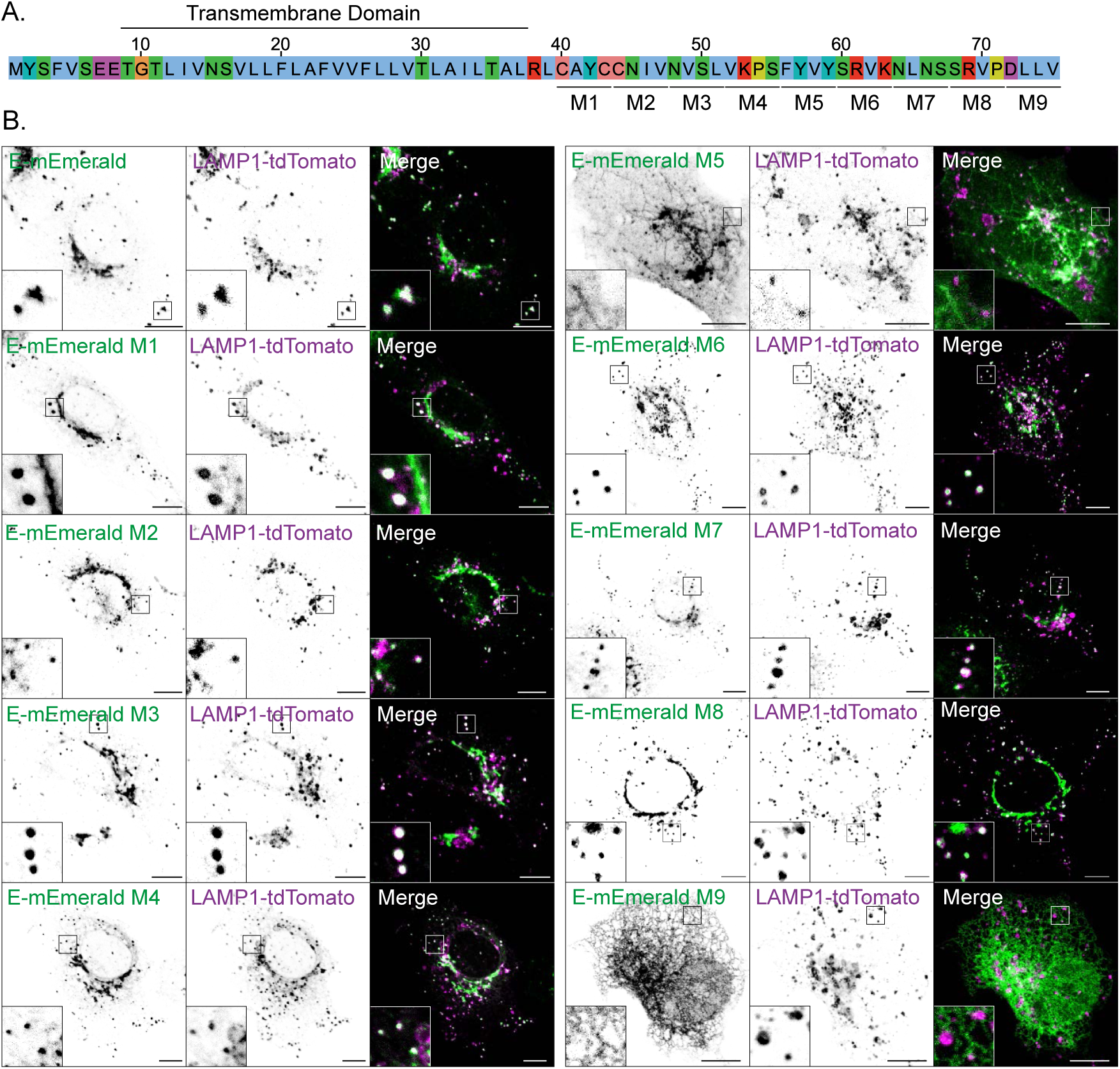
Alanine-scanning mutagenesis of the cytoplasmic tail of E. (**A**) Schematic of alanine-scanning mutagenesis of the C-terminus of E. In mutants M1 – M9, this indicated amino acids were exchanged for Alanine. (**B**) Representative images of VeroE6 cells expressing the indicated versions of E-mEmerald and LAMP1-tdTomato, 15 imaged cells per condition. Scale bars are 10 μm.

**Figure S3.**
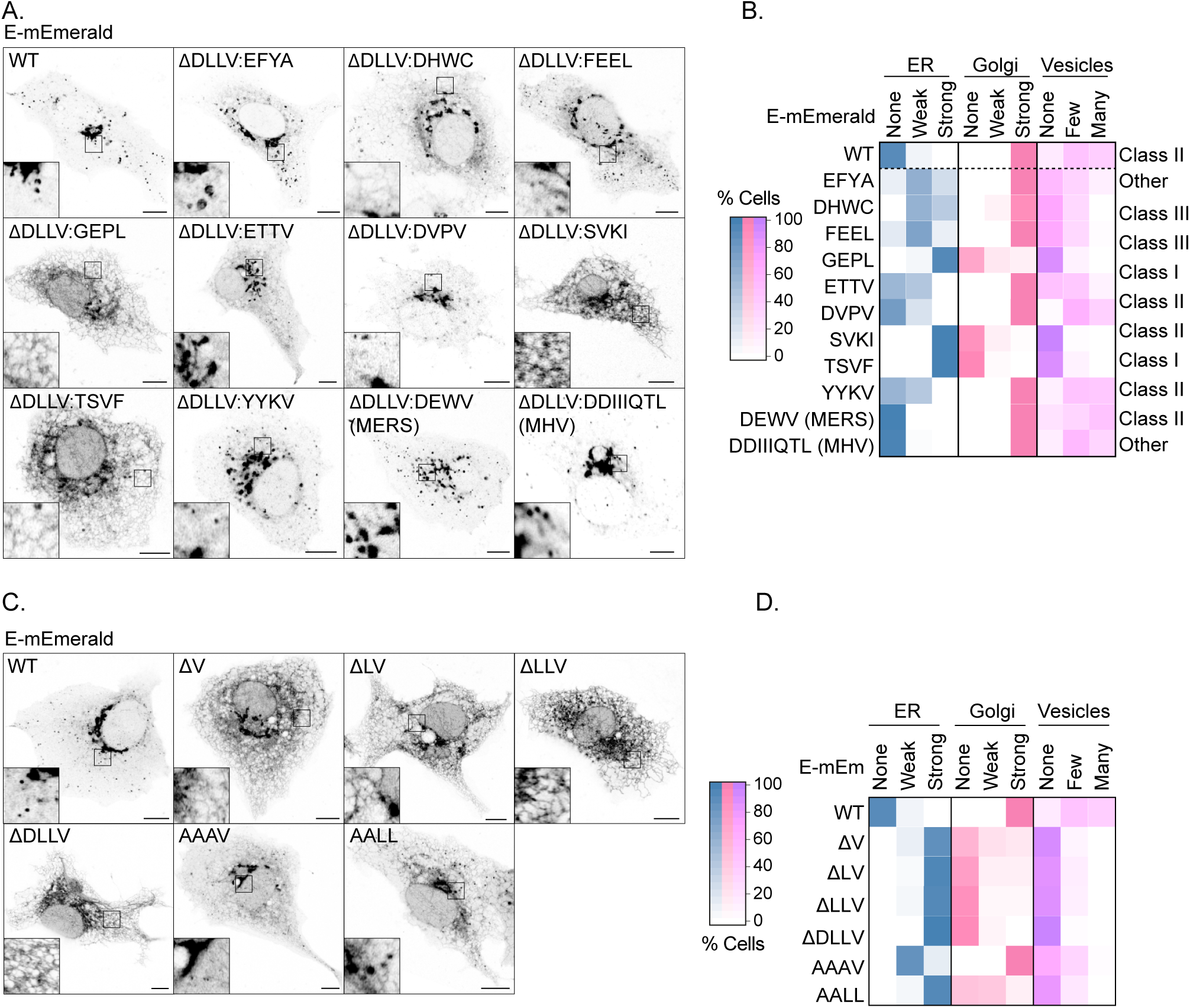
The C-terminus of coronaviral E proteins encodes an ER-export motif. (**A, B**) Representative images and quilt quantification from 50 imaged VeroE6 cells per condition transfected with a panel of E-mEmerald mutants in which the putative C-terminal PDZ-ligand in E (DLLV) was replaced by PDZ-ligands of different classes, or with chimaeric sequences from Middle East Respiratory Virus (MERS)-CoV or Murine Hepatitis Virus (MHV)-CoV (Strain S). The class of PDZ ligand is indicated, with Class I defined by −X-[S/T]-X-ф, Class II defined by - X-ф-X-ф and Class III defined by −X-[D/E/K/R]-X-ф^54^ (**C, D**) Representative images and quilt quantification from 50 imaged VeroE6 cells per condition transfected with a panel of E-mEmerald mutants in which residues in the C-terminal ER-export motif (DLLV) were sequentially deleted or were replaced by C-terminal hydrophobic (AAAV) or di-hydrophobic (AALL) sequences. In microscopy panels, scale bars are 10 μm.

**Figure S4.**
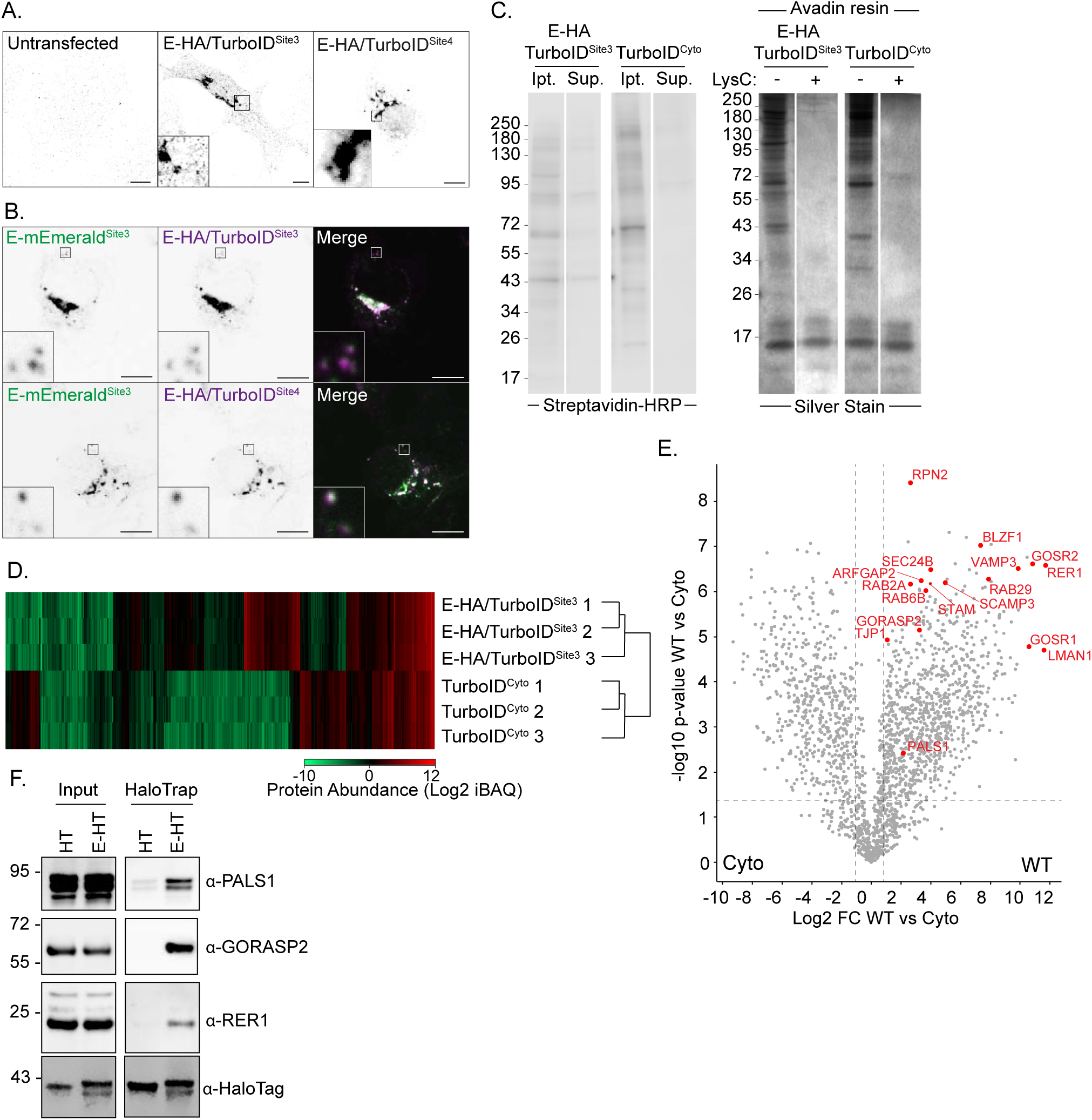
Validation of TurboID-tagging of E proteins. (**A**) VeroE6 cells transfected with plasmids encoding E-HA/TurboID inserted at Site3 or Site4 were fixed and stained with antisera raised against HA. Images representative of 15 or 6 imaged cells respectively. (**B**) 293T cells transfected with plasmids encoding E-HA/TurboID inserted at Site3 or Site4 and E-mEmerald^Site3^ were fixed and stained with antisera raised against HA. Images representative of 5 imaged cells per case. (**C**) Quality control for TurboID-biotinylation-neutravidin pulldown and protein cleavage from beads. Samples (E-HA/TurboID^Site3^ or HA/TurboID (cyto)) were extracted from the post-TurboID reaction pulldown input (Ipt.) and supernatant after neutravidin bead incubation (Sup.), resolved using SDS-PAGE, and blotted using Streptavidin-HRP. Proteins captured on neutravidin beads before and after LysC cleavage were obtained by boiling neutravidin beads in Lamelli buffer with β-mercaptoethanol, were resolved using SDS-PAGE and detected by Silver Stain. (**D**) Hierarchical clustering performed on the median adjusted IBAQ values of each proteomic sample calculated and plotted by average Euclidean distance. Colour scale indicates protein abundance as measured by log_2_ transformed iBAQ values. (**E**) Volcano plot depicting proteins recovered from neutravidin pulldown from 293T cells expressing HA/TurboID (cyto) or E-HA/TurboID (WT) fusions and subject to a 20-minute biotinylation prior to lysis. Volcano plot constructed from N = 3 biological repeats. A selection of proteins strongly enriched by E-HA/TurboID is displayed in red. Percentages were counted from proteins that changed abundance by more than 2-fold and were statistically significant at p < 0.05, as determined by FDR-corrected two-tailed T-tests. (H) Cell lysates and HaloTrap-captured fractions from 293T cells transfected with the indicated E-HT^Site3^ fusions were resolved by SDS-PAGE and examined by western blotting with the indicated antisera (N = 3).

**Figure S5.**
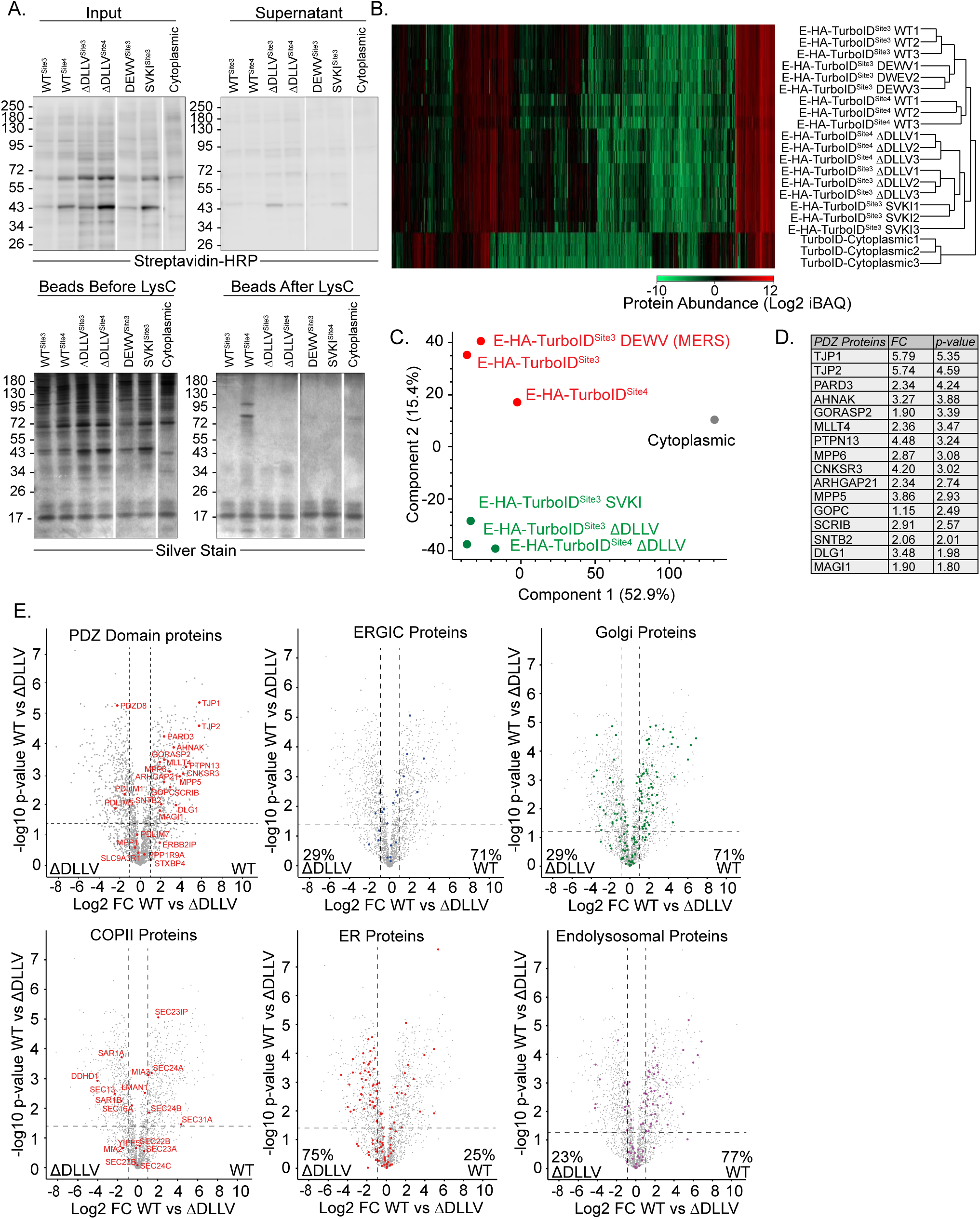
Comparative proximity biotinylation of ER-export proficient and ER-export defective versions of E. (**A**) Quality control for TurboID-biotinylation-neutravidin pulldown and protein cleavage from beads. Samples (E-HA/TurboID^Site3^, E-HA/TurboID^Site4^, E^ΔDLLV^-HA/TurboID^Site3^, E^ΔDLLV^-HA/TurboID^Site4^, E^ΔDLLV+DEWV^-HA/TurboID^Site3^, E^ΔDLLV+SVKI^-HA/TurboID^Site3^, or HA/TurboID (cyto)) were extracted from the post-TurboID reaction pulldown input (Input) and supernatant after neutravidin bead incubation (Supernatant), resolved using SDS-PAGE, and blotted using Streptavidin-HRP. Proteins captured on neutravidin beads before and after LysC cleavage were obtained by boiling neutravidin beads in Lamelli buffer with β-mercaptoethanol, were resolved using SDS-PAGE and detected by Silver Stain. **(B)** Hierarchical clustering performed on the median adjusted IBAQ values of each proteomic sample calculated and plotted by average Euclidean distance. Colour scale indicates protein abundance as measured by log_2_ transformed iBAQ values. (**C**) Principal component analysis (PCA) on median averaged data across the three repeats for each sample. Red dots/text indicate WT-like E proteins, green dots/text indicates ΔDLLV-like E proteins, and grey indicates the cytoplasmic control. (**D**) Tabular depiction of PDZ-domain containing proteins recovered from comparative proteomics of WT and ΔDLLV versions of E-HA/TurboID. (**E**) Volcano plots depicting proteins recovered from a neutravidin pull down from 293T cells expressing either E-HA/TurboID^Site3^ or E-HA/TurboID^Site3^ ΔDLLV, and subject to a 20-minute biotinylation. N = 3. Gene Ontology was used to assign recovered proteins to subcellular localisations. PDZ-domain containing proteins were annotated on the volvano plot. Percentages were counted from proteins that changed abundance by more than 2-fold and were statistically significant at p < 0.05, as determined by FDR-corrected two-tailed T-tests.

**Fig. S6.**
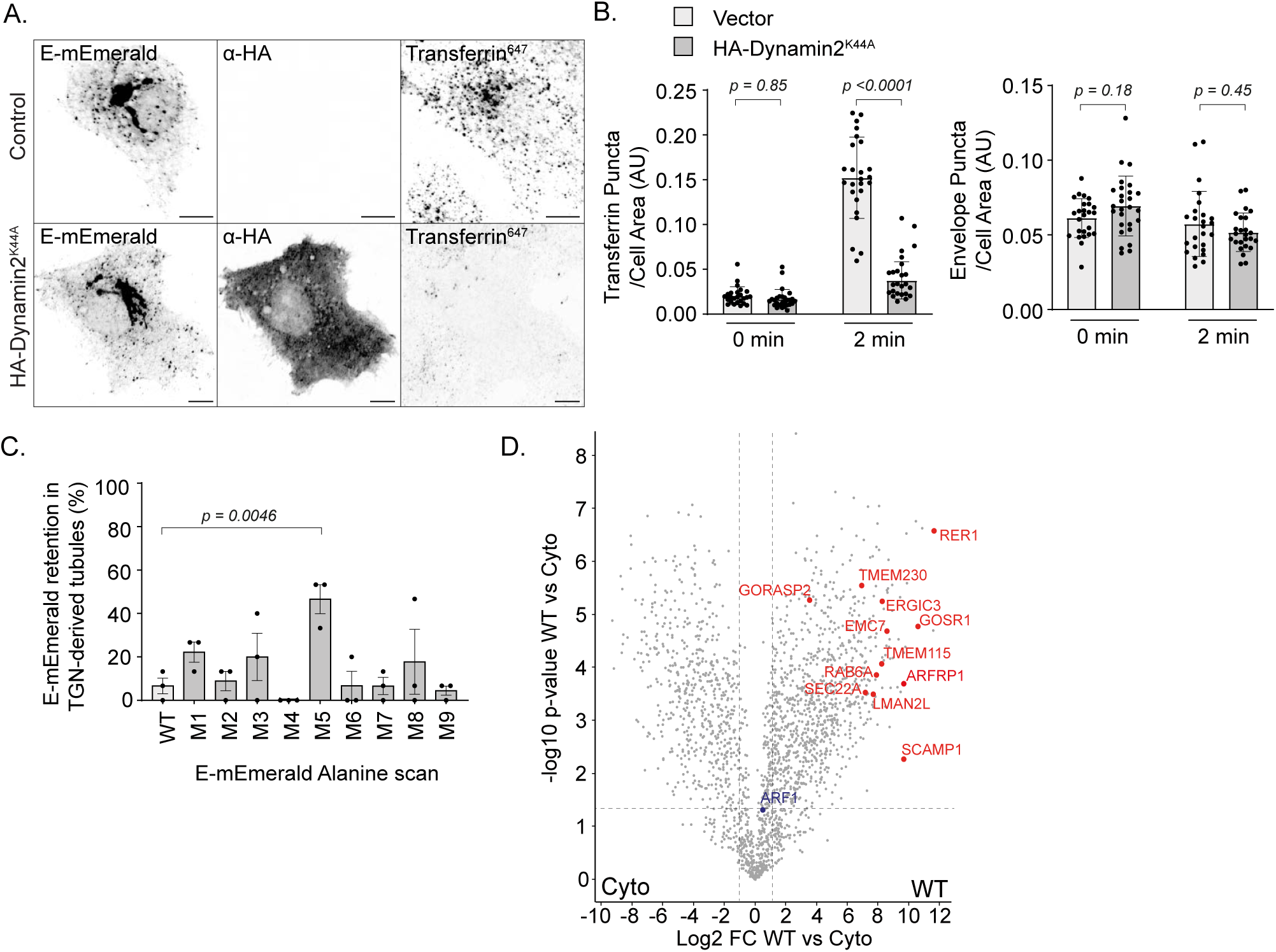
Investigation of Golgi-to-lysosome trafficking of E. (**A**). Representative images of VeroE6 cells transfected for 18 hours with plasmids encoding E-mEmerald and either an empty vector or HA-Dynamin2^K44A^ and then incubated with Alexa^647^-labelled Transferrin for 2 minutes. (**B**) Quantification of Transferrin puncta in cells from A, mean ± S.D. displayed, statistical significance determined by 2-tailed T-Test from 25 imaged cells. Quantification of E-mEmerald puncta in cells from A. Mean ± S.D. displayed with statistical significance determined by 2-tailed T-Test from 25 imaged cells. (**C**) Quantification of E-mEmerald retention of in Golgi-derived tubules in VeroE6 cells transfected with the indicated Alanine-scanning mutants from Figure S2. Quantification is from 15 imaged cells per condition from N = 3 independent experiments. Mean ± S.E. is displayed, with statistical significance determined by one-way ANOVA with each sample compared against WT with Dunnett’s correction applied for multiple testing. (**D**) Volcano plot depicting proteins recovered from a neutravidin pulldown from 293T cells expressing HA/TurboID (cyto) or E-HA/TurboID (WT) fusions and subject to a 20-minute biotinylation prior to lysis. Volcano plot constructed from N = 3 biological repeats. Percentages were counted from proteins that changed abundance by more than 2-fold and were statistically significant at p < 0.05, as determined by FDR-corrected two-tailed T-tests. The position of 12 significantly enriched membrane trafficking proteins selected for our knockout screen are depicted in red. ARF1 is highlighted in blue. In microscopy panels, scale bars are 10 μm.

**Figure S7.**
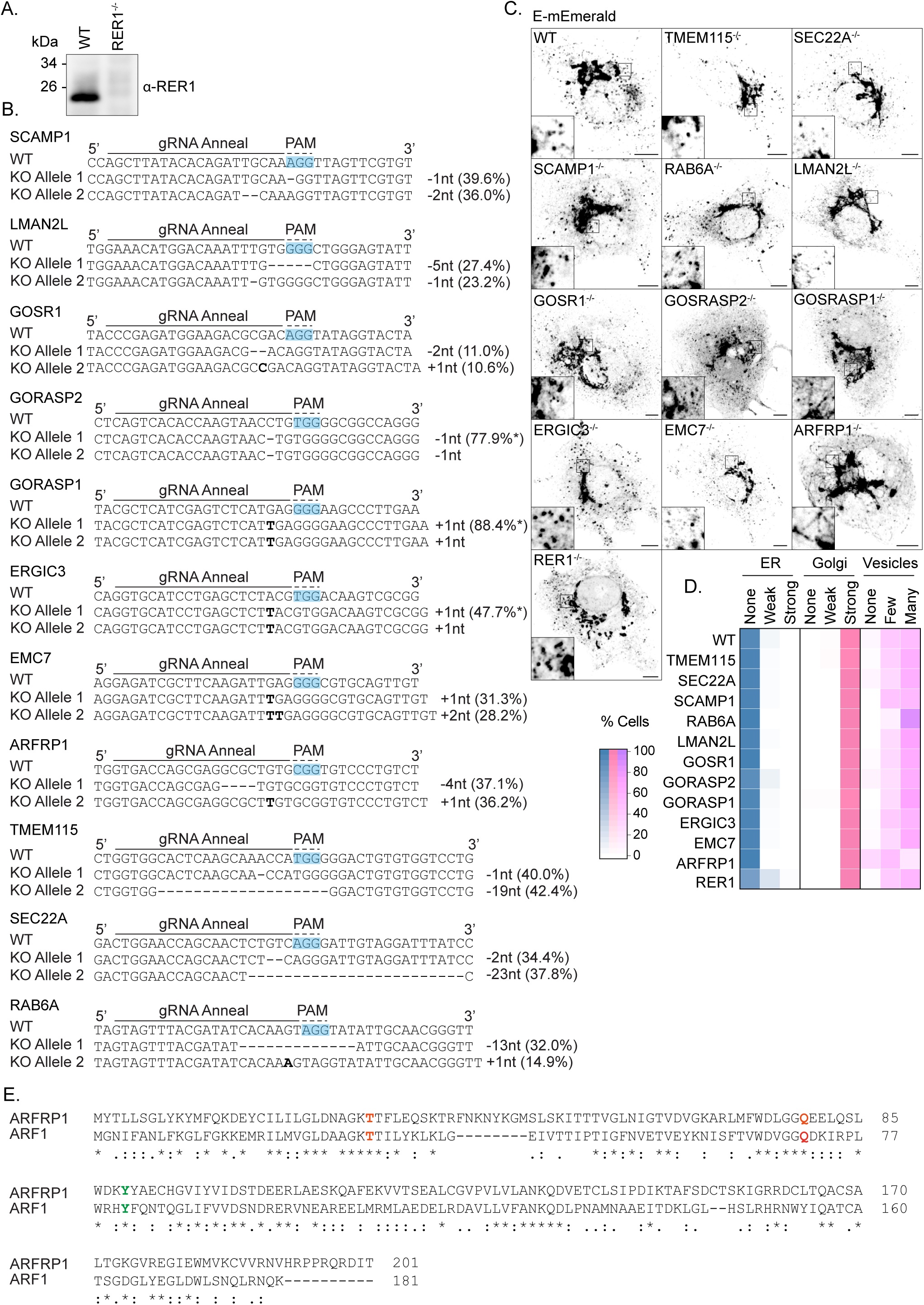
Examination of E-mEmerald trafficking in a panel of knockout VeroE6 cells. (**A**) Resolved cell lysates from WT or RER1^-/-^ VeroE6 cells were examined by western blotting with antisera raised against RER1. (**B**) Genomic locus of the gRNA-annealing region and protospacer adjacent motif (PAM) of each target, and the Next Generation Sequencing result of allele-specific indels in the analysed CRISPR/Cas9-edited VeroE6 clones, including allele percentages of the two most common reads returned. Each edit was verified as a homozygous knockout. (**C, D**) Representative images and quantification of subcellular distribution of the indicated homozygous knockout VeroE6 cells transfected with a plasmid encoding E-mEmerald. Quantification was performed from 50 imaged cells per condition, with localization displayed in a quilt. (**E**) Clustal Omega alignment of ARF1 and ARFRP1 sequences with residues involved in GTPase activity (ARF1: Q71, T31; ARFRP1: Q79, T31) highlighted in red and residues involved in the hydrophobic effector patch (ARF1 Y81; ARFRP1 Y89) highlighted in green.

**Figure S8.**
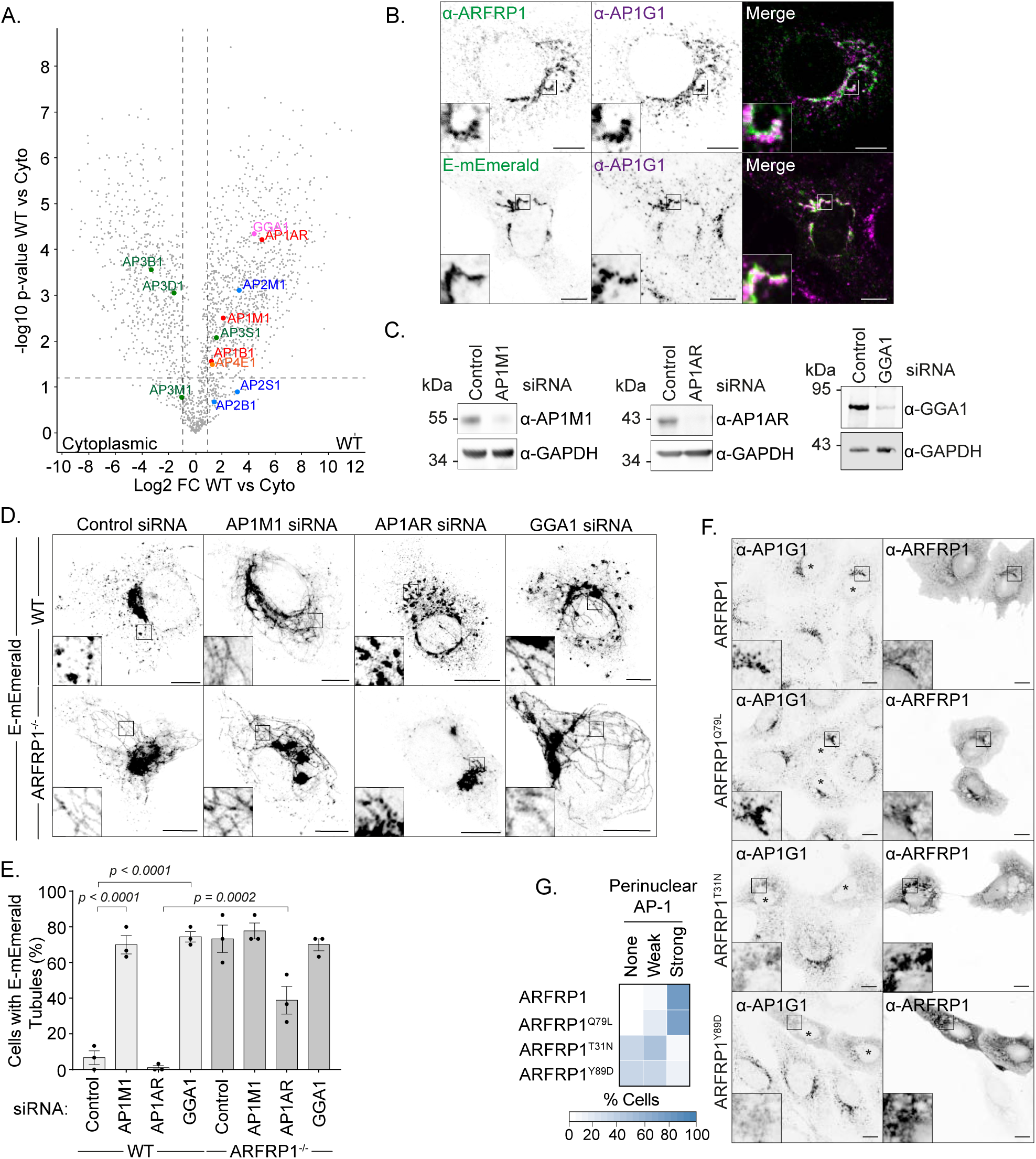
The role of ARFRP1 in coordinating AP-1 at the Golgi for Golgi-to-lysosome trafficking of E. (**A**) Volcano plot depicting proteins recovered from neutravidin pull down from 293T cells expressing either HA-TurboID or E-HA/TurboID and subject to a 20-minute biotinylation. N = 3. All adaptor proteins (AP) identified are highlighted, with colours corresponding to the different AP complexes. See also Fig S6E. Percentages were counted from proteins that changed abundance by more than 2-fold and were statistically significant at p < 0.05, as determined by FDR-corrected two-tailed T-tests. (**B**) VeroE6 cells, or VeroE6 cells transfected with E-mEmerald, were fixed and stained with antisera raised against AP1G1 or ARFRP1 as indicated. (**C**) Resolved cell lysates from VeroE6 cells that had transfected with control siRNA or siRNA targeting AP1M1, AP1AR or GGA1 were examined by western blotting with antisera raised against AP1M1, AP1AR, GGA1 or GAPDH. (**D, E**) Representative images of WT or ARFRP1^-/-^ VeroE6 cells transfected with the indicated siRNA and a plasmid encoding E-mEmerald (D) with quantification of the number of cells displaying E-mEmerald tubules reported (E). 15 cells per experiment were scored from N = 3 independent experiments. Significance calculated by one-way ANOVA comparing WT control against WT AP1M1 siRNA, WT control against GGA1 siRNA, and WT AP1AR siRNA against ARFRP1^-/-^ AP1AR siRNA, with Šidák’s correction for multiple testing. (**F, G**) VeroE6 cells were transfected with plasmids encoding the indicated ARFRP1 proteins and stained with antisera raised against ARFRP1 or AP1G1 (F). Transfected cells indicated by asterisks. Perinuclear localisation of AP1G1 was scored in the accompanying quilt (G) from 50 imaged cells per condition. Acquisition settings were optimised for overexpressed ARFRP1 staining.

**Figure S9.**
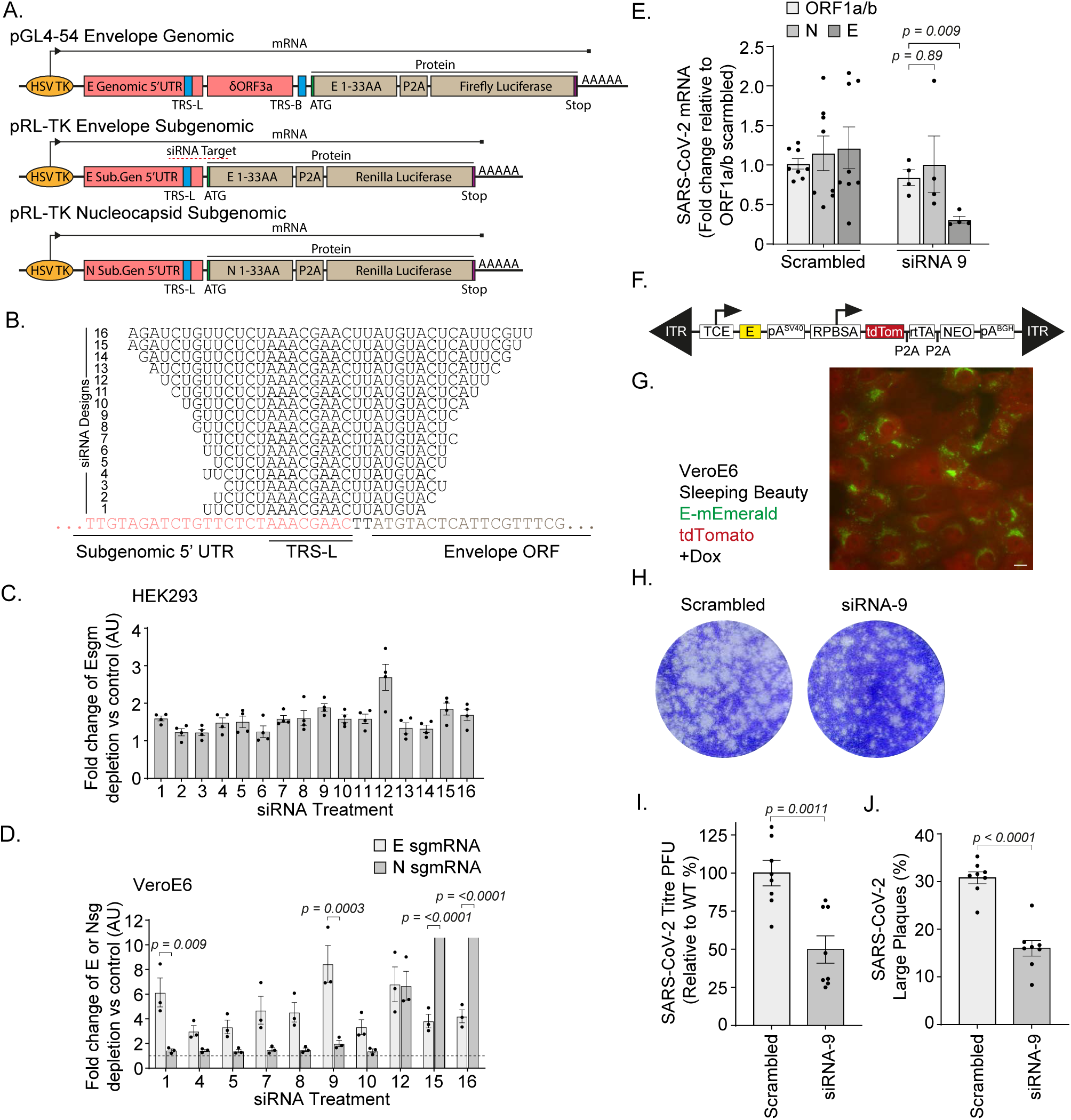
Validation of siRNA designed to target SARS-CoV-2 Envelope sgmRNA. (**A**) Schematic showing the design of a dual-luciferase assay reporter to test siRNA efficacy and specificity against E subgenomic mRNA (sgmRNA). TRS elements and the annealing region of the E-sgmRNAs are labelled. (**B**) Schematic showing the design of E-sgmRNA targeting siRNAs relative to E-sgmRNA. (**C**) For initial screening of E-sgmRNA siRNA potency, Hek293 cells were transfected with 20 µM E-sgmRNA targeting siRNAs or a scrambled siRNA and after 24 hours were transfected with a 1:1 ratio of E-subgenomic/E-genomic dual-luciferase assay reporter plasmids. After 24 hours, luminescence of Firefly and Renilla luciferases was assessed. The ratio of E-sgmRNA reporter and E-genomic reporter was calculated and a fold change relative to scrambled siRNA plotted. Mean ± S.E. is displayed, with N = 3. Relative depletion of E-sgmRNA reported as >1. (**D**) Promising siRNAs from Hek293 cells (C) were tested for efficiency and specificity in VeroE6 cells. VeroE6 were transfected with siRNAs as described above, and cells transfected with either the E-subgenomic/E-genomic dual luciferase reporter, or N-subgenomic/E-genomic dual luciferase reporter. Dual luminescence was recorded as described above. Data plotted shows the fold change of the ratio between E-sgmRNA reporter or N-sgmRNA reporter and E-genomic reporter for E-sgmRNA siRNA treatments, relative to a scrambled control siRNA. Mean ± S.E. is displayed, with N = 3, with statistical testing to test the specificity of the siRNA using a 2-way ANOVA using the Šidák correction for multiple testing. Relative depletion of E-sgmRNA or N-sgmRNA reported as >1. Potent depletion of N-sgmRNA was obtained with siRNA 15 and 16, values for which (15 = 47.54; 16 = 34.86) are omitted from the plot. (**E**) VeroE6 cells were treated with 10 pmol E-sgmRNA targeting siRNA-9, or scrambled siRNA for 20 hours before being infected with SARS-CoV-2 (hCoVL19/England/02/2020) at 1 PFU/cell for 2 hours. After 24 hours, reverse transcription was performed and cDNA levels of SARS-CoV-2 ORF1ab, E and N sgmRNAs were quantified. The data plotted shows viral mRNA expression normalised to actin plotted relative to normalised ORF1a/b expression with the scrambled siRNA treatment. Mean ± S.E. is displayed, with N equal to the number of data points displayed. (**F**) Cartoon of pSBTet-RN. (**G**) VeroE6 cells constitutively expressing tdTomato and expressing a doxycycline-inducible E-mEmerald from the Sleeping Beauty retrotransposition vector were treated with doxycycline and imaged. (**H**) SARS-CoV-2 viruses from VeroE6 cells treated with either scrambled siRNA or E-sgmRNA targeting or siRNA-9 were generated. A diluted stock was applied to confluent VeroE6 cells for 30 minutes. Infected cells were grown for 3 days before being fixed and stained with 0.2% toluidine blue. Images show representative plaque formation. (**I**) Quantification of plaque formation represented as titre (PFU/ml) using the ViralPlaque FIJI macro. Mean ± S.E. presented from N = 8, with significance calculated by a 1-way ANOVA comparing scrambled to E-sgmRNA siRNAs with Dunnet’s correction applied for multiple testing. (**J**) Quantification of the percentage of large plaques *vs* total plaques from plaque assays. Large plaques were defined by having an area greater than 0.82mm^2^ and measured using the ViralPlaque macro in FIJI. Mean ± S.E. presented from N = 8, with significance calculated by a 1-way ANOVA comparing scrambled to E-sgmRNA siRNAs with Dunnet’s correction applied for multiple testing.

## Supplementary Information

**Supplementary Table 1:**

All data from label-free quantification of proximity biotinylation proteomics of HA-TurboID tagged SARS-CoV-2 E, SARS-CoV-2 E mutants, and HA-TurboID cytoplasmic controls.

**Supplementary Table 2:**

Data subset of label free quantification data of proximity biotinylation proteomics of HA-TurboID tagged SARS-CoV-2 E compared to HA-TurboID cytoplasmic control.

**Supplementary Table 3:**

Data subset of label free quantification data of proximity biotinylation proteomics of HA-TurboID tagged SARS-CoV-2 E compared to SARS-CoV-2 E ΔDLLV.

**Supplementary Table 4:**

siRNAs screened for E-sgmRNA targeting.

**Supplementary Table 5:**

PCR primers for the validation of CRISPR knockout clones by next-generation sequencing.

## Experimental Procedures

### Cell Culture

STR-profiled, mycoplasma-free vials of Hek293 (CVCL_0045), 293T (CVCL_0063), VeroE6 cells (CRL-1586, Pasteur) were obtained from the Crick Cell Services Science Technology Platform. Cells were cultured in DMEM containing 10% FBS, Penicillin (100 U/mL) and Streptomycin (0.1 mg/mL).

### Plasmids

Native sequences corresponding to the alpha-variant of SARS-CoV-2 Spike, Nucleocapsid and E cDNAs were purchased from GenScript Biotech: pUC57-2019-NCov-S MC_0101080; pUC57-2019-nCov-N MC_0101085; pUC57-2019-nCOV E MC_0101078. Codon-optimised Spike and Nucleocapsid sequences were kind gifts from Prof. Neil McDonald (Crick) and were cloned similarly into pCR3.1. A sequence corresponding to the native sequence of Membrane was synthesised by GeneWIZ. Coding sequences were amplified by PCR and inserted *EcoRI*-*NotI* into pCR3.1 for mammalian expression. An internal *EcoRI* site in E was removed by silent mutagenesis. Insertion of HaloTag or mEmerald in the E coding sequence was performed using HiFi DNA Assembly, with HaloTag amplified by PCR from pHTN-HaloTag CMV-neo (Promega) and mEmerald amplified by PCR from mEmerald-Sec61b-C1 (Addgene #90992), with a Gly-Gly-Gly-Ser linker placed either side of the HaloTag or mEmerald at Site 3 and Site 4, a single linker placed between E N/C-terminus and HaloTag at tag sites 1 or 5, and a Gly-Gly-Gly-Ser-HaloTag-Gly-Gly-Gly-Ser-Glu-Glu inserted at site 2. Hemagglutinin (HA)-TurboID tagging at sites 3 and 4 was performed by HiFi DNA Assembly, with HA-TurboID amplified by PCR from 3xHA-TurboID-NLS_pCDNA3 (Addgene #107171) and inserted with a Gly-Gly-Gly-Ser linker either side of the HA-TurboID sequence. Emerald-TurboID was used for a cytosolic control in proximity biotinylation experiments and was generated by using HiFi DNA Assembly to assemble Emerald-TurboID in a pLXIN vector, with mEmerald amplified from mEmerald-Sec61b-C1 (Addgene #90992) and TurboID amplified from 3xHA-TurboID-NLS_pCDNA3, with the assembled construct cut with *AgeI* and *EcoRI* to replace EGFP in pEGFP-C1 (Addgene #54759) also digested with *AgeI* and *EcoRI*. pHLARE plasmids were a kind gift from Prof. Diana Barber (University of California, San Francisco). HA-Dynamin2^K44A^ was a kind gift from Prof. Stuart Neil (King’s College London). LAMP1-tdTomato was a kind gift from Dr Max Gutierrez (The Francis Crick Institute). E mutants were generated either by traditional PCR or two-step PCR depending on mutation position. TGN46-mCherry was expressed from pLVX TGN46-mCherry, a kind gift from Prof. David Stephens (University of Bristol, Bristol). TGN46-EGFP was a kind gift from Dr Sharon Tooze (The Francis Crick Institute). GFP controls were expressed from a pCR3.1 GFP-*EcoRI*-*XhoI*-*NotI*^55^. A cDNA encoding mCherry flanked by the 18 amino acid signal sequence from BIP (MKLSLVAAMLLLLSAARA) and a C-terminal KDEL sequence was created by PCR and cloned *EcoRI*-*NotI* into pMSCVneo-*EcoRI*-*XhoI*-*NotI*. A cDNA encoding ARFRP1 was synthesised by GeneWIZ and cloned *EcoRI/NotI* into pCR3.1. Mutations in pCR3.1 ARFRP1 were generated using PCR. An Image clone (OHS5894, clone ID 100000476) encoding human AP1B1 was purchased from Horizon Discovery and the coding sequence was cloned *EcoRI*/*NotI* into pCR3.1 HA-*EcoRI*-*XhoI*-*NotI* using PCR. Envelope 56-63 CoV chimera geneblocks with *EcoRI* and *NotI* overhangs were synthesised by GeneWIZ and cloned into pCR3.1 using the *EcoRI* and *NotI* digestion cloning described above, with mEmerald inserted into site3 as previously described. CRISPR knockouts were performed by transfection with a modified version of px330 (Addgene #42230) which encodes Sniper Cas9^56^ and EBFP2 joined by a P2A site in place of px330’s original Cas9. *BbsI* sites in Sniper-Cas9 (Addgene #42230) were removed by silent mutagenesis, and the ‘px330-Sniper-P2A-BFP’ plasmid was created by HiFI DNA Assembly, with the px330 plasmid linerised to remove Cas9 by PCR, EBFP2 amplified by PCR from mTagBFP2-C1 plasmid (Addgene #54665), *BbsI*-silenced Sniper-Cas9 amplified by PCR, and the P2A synthesised by Integrated DNA Technologies (IDT). To generate px330-Sniper-P2A-BFP plasmids for CRISPR knockouts specific for each gene, overlapping oligonucleotides encoding the gRNA sequence on both parallel and antiparallel strands with *BbsI* compatible overhangs were synthesied by IDT, annealed, and ligated into px330-Sniper-P2A-BFP digested with *BbsI*. Optimal gRNA designs were selected using CRISPick (Broad Institute)^57^. For the dual-luciferase reporter for sgmRNA specificity, pRL-TK Envelope sgmRNA, pRL-TK Nucleocaspid sgmRNA, and pGL4-54 Envelope genomic RNA plasmids were used to assess the effectiveness and specificity of E-sgmRNA targeting siRNAs. pRL-TK plasmids expressed the 5’UTR of E or N sgmRNA including the TRS-L element, the first 99 nucleotides of the CoV-2 protein from the SARS-CoV-2 genomic sequence, and an in-frame P2A linking to Renilla Luciferase. pGL-54 genomic Envelope plasmids expressed the 5’UTR of Envelope genomic RNA including the TRL-B element, δORF3a, the first 99 nucleotides of Envelope from the SARS-CoV-2 Envelope genomic sequence, and an in-frame P2A sequence linking to Firefly Luciferase. pRL-TK plasmids were constructed by digesting the vector with *NheI* and *HindIII* and ligation of the 5’mRNAUTR-99ntORF-P2A insert by HiFi DNA Assembly. The pGL4-54 genomic E plasmid was constructed by digesting the vector with *HindIII*, dephosphorylating using Quick CIP (NEB), and ligation of the 5’genomicUTR-99ntORF-P2A by HiFi DNA Assembly. pRL-TK and pGL4-54 plasmids were from Promega and inserts synthesised by Eurofins. Sleeping Beauty pSBtet-RN^58^ was a kind gift from Dr David Bauer (The Francis Crick Institute). To clone E-Emerald, Emerald, or codon optimised versions of E mutants, Sleeping Beauty vectors were linearised using PCR at the DR insertion sites and inserts encoding these proteins were synthesised (Eurofins), and the plasmids assembled using HiFi DNA assembly. SuperPiggyBack hypertransposase was a kind gift from Prof. Adrian Isaacs (UCL).

### Antibodies and fluorescent labels

An antibody against GAPDH (MAB374) was from Millipore; an antibody against SARS-CoV-2 Spike (GTX632604) was from GeneTex; an antibody against SARS-CoV-2 Nucleocapsid (BS-41408R) was from Bioss; an antibody against SARS-CoV Membrane (101-401-A55) was from (Rockland); an antibody against ERGIC53 (E1031) was from Sigma-Aldrich; an antibody against GM130 (610822) was from BD Biosciences; an antibody against TGN46 (ab50595) was from Abcam; an antibody against EEA1 (610457) was from BD Biosciences; an antibody against HA.11 (16B12) was from Biolegend; an antibody against HaloTag (G9211) was from Promega; an antibody against GFP (7.1/13.1) was from Roche; an antibody against RER1 (HPA051400) was from Sigma-Aldrich; an antibody against PALS1 (17710-1-AP) was from Proteintech; an antibody against GORAPS2 (10598-1-AP) was from Proteintech; an antibody against ARFRP1 (PA5-50606) was from Invitrogen; an antibody against AP1B1 (16932-1-AP) was from Proteintech; an antibody against AP1G (A4200, clone 100/3) was from Sigma; an antibody against GGA1 (25674-1-AP) was from Proteintech; an antibody against AP1AR (NBP1-90879) was from Novus Biologicals; an antibody against AnnexinV conjugated to APC (640932) was from Biolegend; HRP-conjugated Streptavidin (S911) was from Invitrogen; Alexa conjugated secondary antibodies were from Invitrogen and HRP-conjugated secondary antibodies were from Millipore. IRDye 800 CW (925-32210) and IRDye 680 RD (925-68071) were from LI-COR Biosciences. Alexa-647 conjugated Transferrin was from Molecular Probes. Janelia Fluor 646 HaloTag ligand (GA1120), Oregon Green Halotag ligand (G2801), and Tetramethylrhodamine HaloTag ligand (G8251) were from Promega.

### Transient transfection of cDNA

VeroE6 cells were transfected using Lipofectamine-3000 (Life Technologies) according to the manufacturer’s instructions. 293T cells were transfected using linear 25-kDa polyethylenimine (PEI, Polysciences, Inc.), as described previously^59^.

### siRNA Depletion

All siRNA-based depletions were performed at 20 nM final concentration using Lipofectamine RNAiMAX (Life Technologies) transfection reagent according to the manufacturer’s instructions. AP1M1 (L-013196-00-0005), AP1AR (L-015504-02-0005), and GGA1 (M-013694-01-0005) were depleted using ON-TargetPlus SMARTpool siRNAs (Horizon Discovery). A range of custom-made siRNAs (Supplementary Table 4) were synthesised by Horizon Discovery to specifically deplete SARS-CoV-2 E subgenomic mRNA (sg-mRNA). These siRNAs were of different lengths and spanned E’s 5’UTR and ORF.

### Fixed cell imaging

VeroE6 cells were plated at 40,000 per well on 13 mm No. 1.5 coverslips and transfected as described the following day. If cells were transfected with HaloTag-versions of SARS-CoV-2 E, cells were treated with 1 µM Oregon Green Halo ligand for 20 minutes and then washed 3 times with complete media, with a 5 to 10-minute incubation on the final wash. All cells were washed once with PBS before being fixed using 4 % paraformaldehyde for 20 minutes. Cells that required immunolabelling were permeabilised with 0.1 % Triton-X100 in PBS, washed 3 times in PBS, and blocked in 5% FBS for 1 hour. After primary and secondary antibody incubations, coverslips were mounted on X50 SuperFrost microscope slides using Mowiol. Imaging was performed either using a Zeiss LSM 880 as described below, or an Andor Dragonfly 200 spinning disc confocal paired with a Zyla 5.5 sCMOS camera and using a Nikon Eclipse Ti2 with Plan Apo 60x/1.4NA or 100x/1.45NA objectives. To limit overexpression, cells were fixed or imaged 16-18 hours post transfection. E phenotype quantification was performed on 50 cells per condition, with sample identification randomised and blinded during scoring.

### Live cell imaging

Cells stably expressing the indicated proteins, or edited to express fluorescent proteins, were plated in 4- or 8-chamberslides (Ibidi). VeroE6 cells were plated at 40,000 per well in µ-slide ibiTreat 4 well Ibidi chambers and transfected as described the following day. After 16-18 hours, if required, cells were treated with 200 nM JF646 Halo-ligand in complete media for 20 minutes and were then washed twice in growth media before being imaged in FluoroBright DMEM supplemented with 10% FBS, 4 mM L-glutamine, Penicillin (100 U/mL) and Streptomycin (0.1 mg/mL). Airyscan imaging was performed using a Zeiss LSM 880 inverted microscope with a Plan Apo 63X/1.4NA objective fitted with a Fast Live Cell Airyscan detector, definite focus, and heat and CO_2_ incubation. Acquired images were processed using Zeiss’ “Auto” 2D Airyscan processing, and image brightness levels and image crops were adjusted and performed using the FIJI distribution of ImageJ. To limit overexpression, cells were imaged 16 - 18 hours post transfection.

### FLIM-FRET imaging

VeroE6 cells were transfected as previously described with E-mEmerald as the fluorescence donor, and either empty vector for a single colour control or E-HT mutant illuminated with Tetramethylrhodamine (TMR) HaloTag ligand for the fluorescence acceptor. A 1:3 ratio was used for fluorescence donor to acceptor. 5 µM TMR HaloTag ligand was applied to cells for 30 minutes before the cells were washed three times in growth media and then incubated for 30 minutes before cells were imaged in live cell imaging media. FLIM imaging was performed on a Leica TCS SP8 Multiphoton FALCON with a HC PL APO CS2 63x/1.40 oil objective using 470 nm and 552 nm laser lines at 100 Hz scan speed scanning by line, with samples incubated at 37 °C and in 5 % CO_2_. Time-correlated single photon counting fluorescence lifetime data was acquired using a PicoQuant PDL 800-D unit. Raw files were then exported into FLIMfit software^60^ for analysis. Intensity images for each FLIM image were exported into FIJI and a custom script was written to segment each lysosome allowing the fluorescence lifetime of each lysosome to be calculated individually in FLIMfit. The Golgi was excluded from this analysis. For all FLIM analysis, no binning was used, and a single exponential curve was fitted to the data to calculate fluorescence lifetime on a pixel-wise basis.

### Transferrin internalisation assay

VeroE6 cells plated on coverslips were transfected with equivalent amounts of mEmerald tagged E and either empty plasmid or dominant-negative HA-Dynamin2^K44A^. After 16 hours, cells were treated with Transferrin-647 (T23366) purchased from ThermoFisher at 10 µg/mL resuspended in growth media for 2 minutes before being washed once in ice-cold PBS and fixed immediately in 4 % PFA. Untreated cells (0 minutes) were fixed without being treated with Transferrin-647. Cells were prepared for fixed cell imaging, with the presence of HA-Dynamin2^K44A^ detected by detection by an HA antibody. The number of transferrin and E puncta was analysed in FIJI using a custom written script to isolate the lysosomal puncta, with these counts corrected for differing cell area.

### ‘*Quilt*’ Quantification of subcellular distribution

A heatmap-based approach was devised to provide a quickly interpretable graphical display of subcellular localisation across multiple organelles in imaging datasets (*Qu*antitative *I*maging-based *L*ocalisation *T*able). This approach was used to highlight the variability and dominant distributions of E in the secretory pathway and score the perinuclear distribution of AP1 but could be adapted for other classifications of subcellular distribution. To generate a *quilt*, each imaged cell (>50) was scored for strength of localisation of E in the ER, the Golgi, in punctae, at the plasma membrane (not shown), or having a perinuclear distribution in the case of AP-1. For ER, Golgi, and perinuclear AP-1 distributions, ‘None’ was defined by fluorescent images having no reticular or Golgi/AP-1 perinuclear pattern visible, ‘Few’ defined by only a minority of signal being reticular ER or Golgi/AP1 perinuclear relative to rest of the distribution of the fluorescence, and ‘Strong’ defined by the reticular ER or perinuclear Golgi/AP-1 fluorescence being highest or equal highest fluorescence in the image. For punctate localisation, these were scored as ‘None’, ‘Few’, ‘Many’ with this quantification judged subjectively relative to the total cell size. ‘Few’ typically corresponded to <0.045 puncta/µm^2^ and ‘Many’ typically corresponded to >0.045 puncta/µm^2^. No distinction was made between different types of puncta. Golgi was defined by perinuclear fluorescence, and the ER defined by a reticular morphology and nuclear envelope localisation. Once scored, the totals for each compartment category were summated across all the cells imaged in each condition, converted to a percentage of the total number of cells in that condition, and were plotted as a heatmap using R.

### Apoptosis assay

Apoptosis was analysed using the Biolegend APC Annexin V Apoptosis Detection Kit with PI (640932). Cells were plated in 24-well plates at 60,000 cells/well and transfected the following morning with either E-mEmerald^site3^ or GFP. After either 16 hours, 24 hours, 48 hours, or 72 hours, cells were trypsinised, neutralised in complete media, centrifuged, and washed twice in 5 mL of PBS. After the second wash, cell pellets were resuspended in 100 μL of AnnexinV binding buffer and mixed with 5 μL AnnexinV-APC antibody and 10 μL propidium iodide (PI) and incubated at room temperature for 15 minutes. After this time, 400 μL of Annexin V binding buffer was added, and the samples analysed on a Beckman Coulter CytoFLEX LX flow cytometer. GFP/Emerald positive cells were detected using a 488 nm laser and a 525-40 bandpass filter, AnnexinV-APC detected using a 638 nm laser and a 660-10 bandpass filter, and PI detected using a blue laser and a 690-50 bandpass filter. At least 5000 GFP/Emerald events were acquired per sample, and the percentage of GFP+ AnnexinV-APC+ events reported discounting events that were strongly PI+ as dead cells.

### pHLARE Assay

VeroE6 cells were transfected with E-HT and pHLARE plasmids at equal amounts, and after 16 hours cells were incubated with JF646 Halo ligand and imaged by live-cell microscopy on a Zeiss LSM 880 inverted microscope with green, red and far-red channels scanning by line. Images were analysed in FIJI using a custom written script that removed background from all channels, identified lysosomes by their presence in the mCherry red channel, classified these as either E-high or E-low/absent, and then measured the Integrated Density (IntDen) in the 488 channel and the 561 channels. E-high/low lysosomes were determined by eye and equated to at least a 10-fold difference between the mean of the mean grey intensities of the E-high and E-low lysosomes. The ratios between the 488 and 561 channels for E-high and E-low/absent for each lysosome were then averaged across the cell and averaged across the total number of cells recorded. For controls, VeroE6 cells were transfected with pHLARE plasmids alone and then after 16 hours were treated with either 200 nM BafilomycinA1 (19-148) from Sigma-Aldrich for 150 minutes, 10 µM Chloroquine diphosphate (C6628) from Sigma-Aldrich for 160 minutes, 10 mM Ammonium Chloride (254134) from Sigma-Aldrich for 180 minutes, or were untreated, and imaged in green and red channels using the system described. Images were analysed in FIJI using a custom-written script that removed background in all channels and then identified the lysosomes by their presence in the mCherry red channel, and then measured the Integrated Density of each of these lysosomes in both green and red channels. Data was collated as described above for E-HT experiments.

### 4-PBA assay

VeroE6 cells were transfected with Halo tagged SARS-CoV-2 E plasmids for 16 hours. After 16 hours, cells were washed with PBS, and incubated with 1 µM Oregon Green Halo ligand for 20 minutes and washed six times in media, before being incubated with either H_2_O (‘vehicle’), 5 mM, 10 mM, or 20 mM sodium phenylbutyrate (4-PBA) dissolved in H_2_O for 6 hours. After 6 hours, media containing 5 µM TMR Halo ligand supplemented with the appropriate concentration of 4-PBA were added to the cells for 20 minutes. The cells were washed 3 times in media before being fixed and prepared for imaging (as previously described). Sodium phenylbutyrate was from Sigma-Aldrich (SML0309).

### TurboID proximity biotinylation, capture, and mass spectrometry

HEK293T cells were grown for 5 days in biotin-free growth media to remove all sources of biotin. These were then transferred to T75 flasks, with 3 flasks seeded per condition. Cells were transfected with either WT or mutant E-HA/TurboID or E-TurboID (cytosolic control) constructs using PEI, with 1,800 μL optiMEM, 36 μg DNA, and 72 μL PEI used per flask. After exactly 18 hours, cells were biotinylated by incubation in biotin-free growth media supplemented with 50 µM biotin (Sigma-Aldrich). After exactly 20 minutes, flasks were placed on ice and washed once with ice-cold PBS to halt the biotinylation reaction. Cells were scrapped into 10 mL of ice-cold PBS and pelleted, with pellets kept on ice until all samples had been prepared. Cell pellets were lysed by 30 min incubation at 4 °C in 1 mL RIPA buffer (150 mM NaCl, 50 mM Tris-HCl pH8, 1 % NP40, 0.5 % sodium deoxycholate, 0.4 % SDS, 1 mM EDTA) supplemented with cOmplete EDTA-free protease inhibitors (Roche) and 167 U/mL of Benzonase Nuclease (Sigma-Aldrich). During this time pre-acetylated NeutrAvidin agarose beads (Pierce) were washed 4 times with 10 x their volume in lysis buffer, with 40 µL beads used per sample. NeutrAvidin bead acetylation was necessary to stop the Neutravidin being cleaved from the agarose beads during on-bead digestion and performed prior to the day of pulldown by two 30 min incubations of beads with 10 mM Sulfo-NHS acetate (ThermoFisher) on a rotating wheel followed by quenching in 90 mM Tris-HCl pH7.5. Lysed cell pellets were centrifuged at 28,000 x g at 4 °C for 15 minutes to sediment undigested nuclear debris, and supernatants were mixed with equal amounts of washed acetylated NeutrAvidin beads and rotated at room temperature for 2 hours. The beads were then washed 3 times in 500 μL RIPA buffer and 6 times in 25 mM HEPES pH 8.5, with the beads rotated for 3 minutes at room temperature for each wash. After the final wash, beads were resuspended in 100 μL 25 mM HEPES pH 8.5 and 100 ng of Lysyl endopeptidase LysC (WAKO) was added to each sample, with this mixture incubated for 16 hours at 37 °C in a hooded ThermoMixer at 1,200 rpm. Each bead supernatant was then transferred to a new Eppendorf and mixed with 100 ng Trypsin (Pierce) and incubated at 37 °C for 6 hours. The solutions were then acidified to a final concentration of 0.5% trifluoroacetic acid (TFA). Digested samples were loaded onto Evotips and washed once with aqueous acidic buffer (0.1% formic acid in water) before loading onto an Evosep One system coupled to an Orbitrap Fusion Lumos (ThermoFisher Scientific). The Evosep One was fitted with a 15 cm column (PepSep) and a predefined gradient for a 44-minute method was employed. The Orbitrap Lumos was operated in data-dependent mode (1 second cycle time), acquiring IT HCD MS/MS scans in rapid mode after an OT MS1 survey scan (R = 60,000). The MS1 target was 4E5 ions whereas the MS2 target was 1E4 ions. The maximum ion injection time utilized for MS2 scans was 300 ms, the HCD normalized collision energy was set at 32 and the dynamic exclusion was set at 15 seconds. Acquired raw files were processed with MaxQuant v1.5.2.8^61^. Peptides were identified from the MS/MS spectra searched against *Homo sapiens* and SARS-CoV-2 proteomes (UniProt) as well as *Gallus gallus* Avidin (UniProt) and sequences of all TurboID-tagged constructs using Andromeda^62^ search engine. Methionine oxidation, Acetyl (N-term), Acetyl (K) and Deamidation (NQ) were selected as variable modifications. The enzyme specificity was set to Trypsin with a maximum of 2 missed cleavages. The precursor mass tolerance was set to 20 ppm for the first search (used for mass re-calibration) and to 4.5 ppm for the main search. The datasets were filtered on posterior error probability (PEP) to achieve a 1% false discovery rate on protein, peptide and site level. Other parameters were used as pre-set in the software. ‘Unique and razor peptides’ mode was selected to allow identification and quantification of proteins in groups (razor peptides are uniquely assigned to protein groups and not to individual proteins). Intensity based absolute quantification (iBAQ) in MaxQuant was performed using a built-in quantification algorithm^61^ enabling the ‘Match between runs’ option (time window 0.7 minutes) within replicates. MaxQuant output files were processed with Perseus, v1.4.0.2^63^. Data were filtered to remove contaminants, protein IDs originating from reverse decoy sequences and only identified by site. iBAQ intensities were log2 transformed, normalized by median subtraction, and filtered for the presence of 15 valid values. Missing values were imputed from normal distributions. P-values were calculated by two-sample t-tests using Benjamini-Hochberg FDR correction for multiple testing. Crapome data was obtained from Crapome V2^64^. For quality control, four aliquots were taken during sample processing for each sample: ‘Input’, 50 µL of lysate before mixing with beads; ‘Supernatant’, 50 µL of lysate after pull down and pelleting beads; ‘LysC -’, 10 µL of a 100 µL bead resuspension after pull down and bead washing before LysC incubation; ‘LysC +’, 10 µL of a 100 µL bead resuspension after LysC incubation and removal of supernatant containing proteins cleaved from beads. All QC samples were mixed with 4 x LDS with β-ME. ‘Input’ and ‘Supernatant’ QC samples were run on SDS-PAGE gels, blotted, and incubated with Strepatividin-HRP, and ‘LysC -’ and ‘LysC +’ samples were run on SDS-PAGE gels and proteins detected by silver stain using Invitrogen’s SilverQuest staining kit (45-100). All mass spectrometry proteomics data have been deposited to the ProteomeXchange Consortium via the partner repository with the dataset identifier PXD045299. In the depository Site3 and Site 4 are represented by ‘TAL’ and ‘VNVS’ respectively, and the data includes R61A K63A mutants which have been excluded from the final manuscript.

### Immunoprecipitation assay

For verification of proximity biotinylation mass spectrometry data, 25 million 293T cells in a 150 mm dish were transfected with 40 µg pCR3.1 E-HT^Site3^ or pCR3.1 using Polyethyleneimine. After 48 hours, cells were rinsed briefly in ice cold PBS, were lifted from the dish using a cell scraper, collected by centrifugation at 300 x g and lysed on ice in 1 mL of HNG buffer (50 mM Hepes, pH 7.5, 150 mM NaCl, 1 mM EDTA, 1% glycerol) supplemented with 0.5 % Digitonin, protease inhibitors (Complete mini) and phosphatase inhibitors (PhosStop). Lysis was performed in low-bind microfuge tubes (Eppendorf). Insoluble material was removed by centrifugation at 14,000 x g for 2 minutes and the supernatant was incubated with 50 µl HNG-washed agarose beads (Chromotek) with end-over-end rotation for 15 minutes to capture non-specific binding proteins. Beads were collected by centrifugation and discarded. The supernatant was incubated with 50 µl HNG-washed HaloTrap-agarose beads (Chromotek) with end-over-end rotation for 15 minutes hours to capture specific binding proteins. Beads were washed three times in HNG buffer and were transferred to fresh tubes. Bead-bound proteins were released by boiling in 2 x LDS sample buffer and samples analysed using SDS-PAGE and immunoblotting. To investigate the interaction between Envelope or Envelope mutants and HA-tagged AP1B1, 9 million 293T cells were plated in a T75 and transfected the following morning with either 18 µg pCR3.1 E-mEmerald plasmids or 360 ng pCR3.1 GFP, 17.6 µg pCR3.1 and 4.5 µg pCR3.1 HA-AP1B1 as appropriate. To investigate the interaction between Envelope and endogenous AP1B1, 9 million 293T cells were plated in a T75 and transfected the following morning with either 18 µg pCR3.1 E-mEmerald or 360 ng pCR3.1 GFP and 17.6 µg pCR3.1. For all Envelope-AP1 interaction assays, cells were rinsed briefly in ice cold PBS, lifted using a cell scraper, collected by centrifugation at 300 x g, and lysed on ice in 1 mL of HNG buffer supplemented with 1% NP40, protease inhibitors and phosphatase inhibitors. Lysates were rotated at for 30 minutes at 4 °C and insoluble material was removed by centrifugation at 14,000 x rpm for 15 minutes at 4 °C. Supernatants were added to 15 µl of GFPTrap-magnetic agarose beads (Chromotek) and incubated with end-over-end rotation for 30 minutes at 4 °C to capture specific binding proteins. Beads were washed five times in HNG buffer supplemented with 0.1 % NP40 with 1 minute of end-over-end rotation for each wash. Samples were eluted from beads by incubation at 95 °C with 40 µL 2x LDS sample buffer containing β-mercaptoethanol for 5 minutes and analysed using SDS-PAGE and immunoblotting.

### CRISPR-KO and validation

VeroE6 cells were transfected with px330-Sniper-P2A-BFP plasmids each cloned to express a gRNA specific for each gene of interest. After 2 days, cells were single-cell sorted using a FACSAria Fusion flow cytometer (BD Biosciences) into 96-well plates enriching for BFP-positive populations. 20 clones from each plate were expanded and genomic DNA extracted by a GeneJet Genomic DNA Purification Kit (ThermoFisher Scientific, K0722) following the manufacturer’s protocol. PCR was then used to generate amplicons of between 150-400 nucleotides that surrounded the gRNA annealing region and PAM of the gene of interest, with forward primers designed with a TCGTCGGCAGCGTCAGATGTGTATAAGAGACAG overhang and reverse primers with a GTCTCGTGGGCTCGGAGATGTGTATAAGAGACAG overhang encoding adaptors for Nextera XT Indexing (Illumina). Primers are listed in Supplementary Table 5. PCR products were purified using Ampure XP beads (Beckman Coulter) following the manufacturer’s instrctions, with the size of the product verified by agarose gel electrophoresis. Nextera XT Indexing primers were then used to index amplicons by PCR, and the product purified by Ampure XP beads. Samples were then sequenced using a MiSqeq (Illumina) and alignment of NGS results to the genome was performed using SerialCloner software with allele specific reads reported at approximately a 50:50 ratio. Homozygous RER1 knockout and ARFRP1 knockout was validated by immunoblotting.

### Luciferase Assay for Envelope sg-mRNA siRNA Screening

Luciferase screening of Envelope subgenomic siRNA potentency and specificity was performed using the Dual-Luciferase Reporter Assay System (Promega, E19190). For initial screening of siRNA potency, Hek293 cells were plated in 24-well plates at 50,000 cells per well and after 4 hours were transfected with 20 µM custom E-sgmRNA targeting siRNAs or a scrambled control siRNA. After 24 hours, cells were transfected with a 1:1 ratio of pRL-TK Envelope subgenomic and pGL4-54 Envelope genomic plasmids, with transfection media changed 6 hours post transfection. After 24 hours, cells were washed once in PBS and lysed by addition of 100 µL of 1X Passive Lysis Buffer (Promega) and incubated on a table-top rocker for 15 minutes at room temperature. Lysate was then collected and 20 µL was mixed with 100 µL LARII reagent and Firefly Luciferase luminescence measured by a Promega Glomax 20/20 Luminometer. 100 µL of Stop & Glo reagent was then added and the Renilla Luciferase luminesence measured. siRNA potency was determined by the fold-change increase of the Luciferase:Renilla ratio compared a control of a non-targeting siRNA. The top 10 performing siRNAs were then validated for potency and specificity in VeroE6 cells. VeroE6 cells were plated and treated with siRNA as described above. Cells were then transfected with either a 1:1 ratio of pRL-TK Envelope sgmRNA plasmid and pGL4-54 Envelope genomic plasmid, or pRL-TK Nucleocapsid sgmRNA plasmd and pGL4-54 Envelope genomic plasmid, and measurements of Firefly and Renilla luminescence measured as before. siRNA potency was assessed as described previously, and siRNA specificity determined by the fold-change of the siRNA’s E^genomic^/E^subgenomic^ ratio against a non-targeting control compared to a E^genomic^/N^subgenomic^ ratio against a non-targeting control.

### Viruses and infection

The SARS-CoV-2 isolate used (hCoVL19/England/02/2020) was obtained from the Respiratory Virus Unit, Public Health England, UK. Virus stocks were propagated in Vero V1 cells (a gift from Stephen Goodbourn, St George’s University of London) by infection at an MOI of 0.0001 in DMEM, supplemented with 2 % foetal calf serum and penicillin-streptomycin (100 U/mL each) and harvested after 4 days. Stocks were titrated on Vero E6 cells (Pasteur). Vero cells were transfected in 24-well plates with 10 pmol siRNA using Lipofectamine 3000 (Invitrogen). After 2 hours, the media was replaced with 10 % FCS DMEM containing 0.5 µg/mL doxycycline hydrochloride (ThermoScientific). After 20 hours, cells were infected with SARS-CoV-2 at a multiplicity of 1 PFU/cell, in DMEM containing 2 % FCS and 50 μg/mL DEAE-dextran. After 2 hours, the inoculum was replaced with 2 % FCS DMEM containing 0.5 µg/mL doxycycline hydrochloride and cells were transfected again with 10 pmol siRNAs to ensure maximal knockdown. Cells were incubated at 37 °C for 24 hours, before supernatants were harvested for plaque assay and cells were harvested in Trizol (Invitrogen) for qPCR analysis. For *trans*-complementation assays, bearing the sleeping beauty system, tdTomato-positive cells were obtained by FACS as described below.

### Plaque assay

Confluent VeroE6 cells were infected with diluted supernatants for 30 minutes. Overlay medium (1x MEM, 1.2 % Avicel and 100 U/mL each penicillin-streptomycin) was added and cells were incubated at 37 °C for 3 days. Cells were fixed with 4 % PFA in PBS and stained using 0.2 % toluidine blue (Sigma). Plaque area was determined using the ViralPlaque macro in FIJI^65^. Large plaques were defined by having an area greater than 0.82 mm^2^ and measured using the ViralPlaque macro in FIJI^65^.

### Virus Like Particle production assay

7.5 million 293T cells in a 100 mm dish or T75 were transfected with a mixture comprising 5 µg pCR3.1 SARS-CoV-2 S, 3 µg pCR3.1 SARS-CoV-2 M, 3 µg pCR3.1 SARS-CoV-2 E (or derivatives) and 1 µg of pCR3.1 SARS-CoV-2 N. Codon optimised sequences were used in all cases. Media was changed after 6 hours. 48 hours after transfection, supernatants were clarified by centrifugation (300 x g, 2 minutes) and passed through a 0.45 µm syringe filter. Supernatants were underlaid with a PBS 20% sucrose cushion and subject to ultracentrifugation in a Beckman SW41 Ti swinging bucket rotor at 28,000 rpm for 3 hours at 4 °C. Supernatants were removed and pellets were resuspended in 30 µL PBS and incubated overnight at 4 °C. The next morning, 30 µL of 2 x LDS-sample buffer was added for sample recovery. Cellular fractions were obtained by lifting cells with PBS and collecting them by centrifugation (300 x g, 2 minutes) before resuspending the pellet in fresh PBS and adding an equal volume of 2 x LDS sample buffer.

### RNA extraction and qPCR

RNA was extracted using the Direct-Zol miniprep kit (Zymo). cDNA was synthesised using Superscript VILO Master Mix (Invitrogen). SARS-CoV-2 ORF1ab and Actin were quantified using the 2019-nCoV: Real-Time Fluorescent RT-PCR kit (BGI). Envelope and Nucleocapsid subgenomic mRNAs were quantified using specific primer probe sets from^41^ (sequences below), using Taqman Multiplex Master Mix (Applied Biosystems). Viral gene expression was normalised to Actin expression and expressed as a fold change compared to the scrambled siRNA control cells.

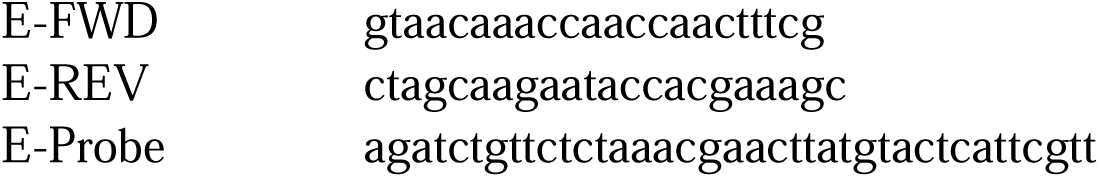
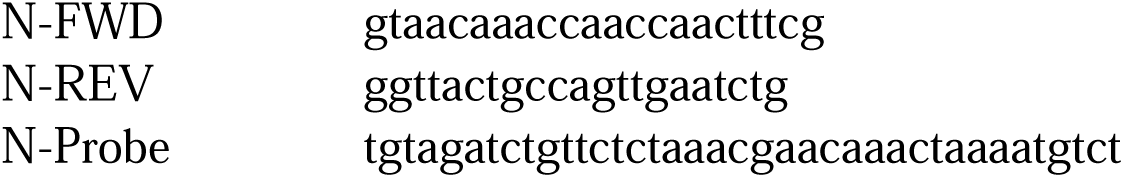

### Sleeping Beauty Generation and Flow Cytometry

Sleeping Beauty plasmids (pSBtet-RN) expressing codon optimised versions of E, E-Emerald or GFP were cloned as previously described and transfected into VeroE6 cells at a 1:1 ratio with the SuperPiggyBac transposase using Lipofectamine 3000. Cells were selected with G418 for 1 week before being grown in normal DMEM media. Due to the low genomic transposition efficiency 3 rounds of flow cytometry enrichment were performed to achieve high proportions of transposed cells. An E-mEmerald line was generated in order to compare the level of constitutive tdTomato expression that corresponded to dox-inducible E or E-mEmerald expression. This was determined by treatment of Sleeping Beauty E-mEmerald cells with and without 0.5 µg/mL doxycycline hydrochloride for 20 hrs and assaying for mEmerald positive cells (detected using a 488 nm laser and a 525-40 bandpass filter) and tdTomato positive cells (detected using a 561 nm laser and a 610-20 bandpass filter. The brightness of the appropriate tdTomato population was determined by using 8-peak fluorescent beads, allowing the brightness of this population to be quantified. Enrichment of tdTomato-positive populations of an equivalent brightness was performed on a BD FACSAria Fusion flow cytometry sorter gating on the appropriate tdTomato positive population as determined by acquisition using a 561 nm laser and a 610-20 bandpass filter and plotting against acquisition using a 488 nm laser and a 530-30 bandpass filter to discount autofluorescence. Three rounds of enrichment sorting were performed to enrich the appropriate tdTomato population to >90% prior to use in infectivity assays.

### Sequence alignments

Alignments of coronavirus Envelope sequences were performed using T-Coffee^66^. Aligned sequences were then exported and viewed in Jalview^67^ and residues colour coded using ClustalX colour map.

### Gene Ontology analysis

Gene ontology cellular compartment (GO:CC) data was used to categorise the subcellular distribution of proteins identified from proteomic analysis. The GO:CC terms used were ER: GO:0005783, ERGIC: GO:0005793, Golgi: GO:0005794, Lysosomal: GO:0005768 (endosome), GO:0005764 (lysosome).

### Statistical analysis

2-tailed Student’s T-tests, or ordinary 1- or 2-way ANOVA with the indicated corrections for multiple testing were used to assess significance between test samples and controls and were performed using GraphPad Prism.

## References

1. Gordon, D.E., Jang, G.M., Bouhaddou, M., Xu, J., Obernier, K., White, K.M., O’Meara, M.J., Rezelj, V.V., Guo, J.Z., Swaney, D.L., et al. (2020). A SARS-CoV-2 protein interaction map reveals targets for drug repurposing. Nature 583, 459–468. 10.1038/s41586-020-2286-9.

2. Stertz, S., Reichelt, M., Spiegel, M., Kuri, T., Martínez-Sobrido, L., García-Sastre, A., Weber, F., and Kochs, G. (2007). The intracellular sites of early replication and budding of SARS-coronavirus. Virology 361, 304–315. 10.1016/j.virol.2006.11.027.

3. Schoeman, D., and Fielding, B.C. (2019). Coronavirus envelope protein: current knowledge. Virol. J. 16, 69. 10.1186/s12985-019-1182-0.

4. Ho, Y., Lin, P.-H., Liu, C.Y.Y., Lee, S.-P., and Chao, Y.-C. (2004). Assembly of human severe acute respiratory syndrome coronavirus-like particles. Biochem. Biophys. Res. Commun. 318, 833–838. 10.1016/j.bbrc.2004.04.111.

5. Siu, Y.L., Teoh, K.T., Lo, J., Chan, C.M., Kien, F., Escriou, N., Tsao, S.W., Nicholls, J.M., Altmeyer, R., Peiris, J.S.M., et al. (2008). The M, E, and N Structural Proteins of the Severe Acute Respiratory Syndrome Coronavirus Are Required for Efficient Assembly, Trafficking, and Release of Virus-Like Particles. J. Virol. 82, 11318–11330. 10.1128/jvi.01052-08.

6. Hsieh, P.-K., Chang, S.C., Huang, C.-C., Lee, T.-T., Hsiao, C.-W., Kou, Y.-H., Chen, I.-Y., Chang, C.-K., Huang, T.-H., and Chang, M.-F. (2005). Assembly of severe acute respiratory syndrome coronavirus RNA packaging signal into virus-like particles is nucleocapsid dependent. J. Virol. 79, 13848–13855. 10.1128/jvi.79.22.13848-13855.2005.

7. Nieto-Torres, J.L., DeDiego, M.L., Álvarez, E., Jiménez-Guardeño, J.M., Regla-Nava, J.A., Llorente, M., Kremer, L., Shuo, S., and Enjuanes, L. (2011). Subcellular location and topology of severe acute respiratory syndrome coronavirus envelope protein. Virology 415, 69–82. 10.1016/j.virol.2011.03.029.

8. Cohen, J.R., Lin, L.D., and Machamer, C.E. (2011). Identification of a Golgi complex-targeting signal in the cytoplasmic tail of the severe acute respiratory syndrome coronavirus envelope protein. J. Virol. 85, 5794–5803. 10.1128/jvi.00060-11.

9. Mandala, V.S., McKay, M.J., Shcherbakov, A.A., Dregni, A.J., Kolocouris, A., and Hong, M. (2020). Structure and drug binding of the SARS-CoV-2 envelope protein transmembrane domain in lipid bilayers. Nat. Struct. Mol. Biol. 27, 1202–1208. 10.1038/s41594-020-00536-8.

10. Xia, B., Shen, X., He, Y., Pan, X., Liu, F.-L., Wang, Y., Yang, F., Fang, S., Wu, Y., Duan, Z., et al. (2021). SARS-CoV-2 envelope protein causes acute respiratory distress syndrome (ARDS)-like pathological damages and constitutes an antiviral target. Cell Res. 31, 847–860. 10.1038/s41422-021-00519-4.

11. Venkatagopalan, P., Daskalova, S.M., Lopez, L.A., Dolezal, K.A., and Hogue, B.G. (2015). Coronavirus envelope (E) protein remains at the site of assembly. Virology 478, 75–85. 10.1016/j.virol.2015.02.005.

12. DeDiego, M.L., Alvarez, E., Almazán, F., Rejas, M.T., Lamirande, E., Roberts, A., Shieh, W.-J., Zaki, S.R., Subbarao, K., and Enjuanes, L. (2006). A severe acute respiratory syndrome coronavirus that lacks the E gene is attenuated in vitro and in vivo. J. Virol. 81, 1701–1713. 10.1128/jvi.01467-06.

13. Kuo, L., and Masters, P.S. (2003). The Small Envelope Protein E Is Not Essential for Murine Coronavirus Replication. J. Virol. 77, 4597–4608. 10.1128/jvi.77.8.4597-4608.2003.

14. Ortego, J., Ceriani, J.E., Patiño, C., Plana, J., and Enjuanes, L. (2007). Absence of E protein arrests transmissible gastroenteritis coronavirus maturation in the secretory pathway. Virology 368, 296–308. 10.1016/j.virol.2007.05.032.

15. Ghosh, S., Dellibovi-Ragheb, T.A., Kerviel, A., Pak, E., Qiu, Q., Fisher, M., Takvorian, P.M., Bleck, C., Hsu, V.W., Fehr, A.R., et al. (2020). β-Coronaviruses Use Lysosomes for Egress Instead of the Biosynthetic Secretory Pathway. Cell 183, 1520–1535.e14. 10.1016/j.cell.2020.10.039.

16. Wang, W.-A., Carreras-Sureda, A., and Demaurex, N. (2023). SARS-CoV-2 infection alkalinizes the ERGIC and lysosomes through the viroporin activity of the viral envelope protein. J. Cell Sci. 136, jcs260685. 10.1242/jcs.260685.

17. Xia, B., Wang, Y., Pan, X., Cheng, X., Ji, H., Zuo, X., Jiang, H., Li, J., and Gao, Z. (2022). Why is the SARS-CoV-2 Omicron variant milder? Innov. 3, 100251. 10.1016/j.xinn.2022.100251.

18. Webb, B.A., Aloisio, F.M., Charafeddine, R.A., Cook, J., Wittmann, T., and Barber, D.L. (2021). pHLARE: a new biosensor reveals decreased lysosome pH in cancer cells. Mol. Biol. Cell 32, 131–142. 10.1091/mbc.e20-06-0383.

19. Jimenez-Guardeño, J.M., Nieto-Torres, J.L., DeDiego, M.L., Regla-Nava, J.A., Fernandez-Delgado, R., Castaño-Rodriguez, C., and Enjuanes, L. (2014). The PDZ-Binding Motif of Severe Acute Respiratory Syndrome Coronavirus Envelope Protein Is a Determinant of Viral Pathogenesis. PLoS Pathog. 10, e1004320. 10.1371/journal.ppat.1004320.

20. Nufer, O., Guldbrandsen, S., Degen, M., Kappeler, F., Paccaud, J.-P., Tani, K., and Hauri, H.-P. (2002). Role of cytoplasmic C-terminal amino acids of membrane proteins in ER export. J. Cell Sci. 115, 619–628. 10.1242/jcs.115.3.619.

21. Ma, W., Goldberg, E., and Goldberg, J. (2017). ER retention is imposed by COPII protein sorting and attenuated by 4-phenylbutyrate. eLife 6, e26624. 10.7554/elife.26624.

22. Branon, T.C., Bosch, J.A., Sanchez, A.D., Udeshi, N.D., Svinkina, T., Carr, S.A., Feldman, J.L., Perrimon, N., and Ting, A.Y. (2018). Efficient proximity labeling in living cells and organisms with TurboID. Nat. Biotechnol. 36, 880–887. 10.1038/nbt.4201.

23. Chai, J., Cai, Y., Pang, C., Wang, L., McSweeney, S., Shanklin, J., and Liu, Q. (2021). Structural basis for SARS-CoV-2 envelope protein recognition of human cell junction protein PALS1. Nat. Commun. 12, 3433. 10.1038/s41467-021-23533-x.

24. Teoh, K.-T., Siu, Y.-L., Chan, W.-L., Schlüter, M.A., Liu, C.-J., Peiris, J.S.M., Bruzzone, R., Margolis, B., and Nal, B. (2010). The SARS Coronavirus E Protein Interacts with PALS1 and Alters Tight Junction Formation and Epithelial Morphogenesis. Mol. Biol. Cell 21, 3838–3852. 10.1091/mbc.e10-04-0338.

25. Laulagnier, K., Schieber, N.L., Maritzen, T., Haucke, V., Parton, R.G., and Gruenberg, J. (2011). Role of AP1 and Gadkin in the traffic of secretory endo-lysosomes. Mol. Biol. Cell 22, 2068–2082. 10.1091/mbc.e11-03-0193.

26. Hirst, J., Edgar, J.R., Borner, G.H.H., Li, S., Sahlender, D.A., Antrobus, R., and Robinson, M.S. (2015). Contributions of epsinR and gadkin to clathrin-mediated intracellular trafficking. Mol. Biol. Cell 26, 3085–3103. 10.1091/mbc.e15-04-0245.

27. Schmidt, M.R., Maritzen, T., Kukhtina, V., Higman, V.A., Doglio, L., Barak, N.N., Strauss, H., Oschkinat, H., Dotti, C.G., and Haucke, V. (2009). Regulation of endosomal membrane traffic by a Gadkin/AP-1/kinesin KIF5 complex. Proc. Natl. Acad. Sci. 106, 15344–15349. 10.1073/pnas.0904268106.

28. Buser, D.P., and Spang, A. (2023). Protein sorting from endosomes to the TGN. Front. Cell Dev. Biol. 11, 1140605. 10.3389/fcell.2023.1140605.

29. Antonny, B., Burd, C., Camilli, P.D., Chen, E., Daumke, O., Faelber, K., Ford, M., Frolov, V.A., Frost, A., Hinshaw, J.E., et al. (2016). Membrane fission by dynamin: what we know and what we need to know. EMBO J. 35, 2270–2284. 10.15252/embj.201694613.

30. Chavrier, P., and Ménétrey, J. (2010). Toward a Structural Understanding of Arf Family:Effector Specificity. Structure 18, 1552–1558. 10.1016/j.str.2010.11.004.

31. Rebendenne, A., Roy, P., Bonaventure, B., Valadão, A.L.C., Desmarets, L., Arnaud-Arnould, M., Rouillé, Y., Tauziet, M., Giovannini, D., Touhami, J., et al. (2022). Bidirectional genome-wide CRISPR screens reveal host factors regulating SARS-CoV-2, MERS-CoV and seasonal HCoVs. Nat. Genet. 54, 1090–1102. 10.1038/s41588-022-01110-2.

32. Israeli, M., Finkel, Y., Yahalom-Ronen, Y., Paran, N., Chitlaru, T., Israeli, O., Cohen-Gihon, I., Aftalion, M., Falach, R., Rotem, S., et al. (2022). Genome-wide CRISPR screens identify GATA6 as a proviral host factor for SARS-CoV-2 via modulation of ACE2. Nat. Commun. 13, 2237. 10.1038/s41467-022-29896-z.

33. Biering, S.B., Sarnik, S.A., Wang, E., Zengel, J.R., Leist, S.R., Schäfer, A., Sathyan, V., Hawkins, P., Okuda, K., Tau, C., et al. (2022). Genome-wide bidirectional CRISPR screens identify mucins as host factors modulating SARS-CoV-2 infection. Nat. Genet. 54, 1078–1089. 10.1038/s41588-022-01131-x.

34. Ohno, H., Stewart, J., Fournier, M.-C., Bosshart, H., Rhee, I., Miyatake, S., Saito, T., Gallusser, A., Kirchhausen, T., and Bonifacino, J.S. (1995). Interaction of Tyrosine-Based Sorting Signals with Clathrin-Associated Proteins. Science 269, 1872–1875. 10.1126/science.7569928.

35. Edeling, M.A., Mishra, S.K., Keyel, P.A., Steinhauser, A.L., Collins, B.M., Roth, R., Heuser, J.E., Owen, D.J., and Traub, L.M. (2006). Molecular Switches Involving the AP-2 β2 Appendage Regulate Endocytic Cargo Selection and Clathrin Coat Assembly. Dev. Cell 10, 329–342. 10.1016/j.devcel.2006.01.016.

36. Owen, D.J., and Evans, P.R. (1998). A Structural Explanation for the Recognition of Tyrosine-Based Endocytotic Signals. Science 282, 1327–1332. 10.1126/science.282.5392.1327.

37. Martin-Serrano, J., and Bieniasz, P.D. (2003). A Bipartite Late-Budding Domain in Human Immunodeficiency Virus Type 1. J. Virol. 77, 12373–12377. 10.1128/jvi.77.22.12373-12377.2003.

38. Zhang, X., Liu, Y., Liu, J., Bailey, A.L., Plante, K.S., Plante, J.A., Zou, J., Xia, H., Bopp, N.E., Aguilar, P.V., et al. (2021). A trans-complementation system for SARS-CoV-2 recapitulates authentic viral replication without virulence. Cell 184, 2229–2238.e13. 10.1016/j.cell.2021.02.044.

39. Sola, I., Almazán, F., Zúñiga, S., and Enjuanes, L. (2015). Continuous and Discontinuous RNA Synthesis in Coronaviruses. Annu. Rev. Virol. 2, 265–288. 10.1146/annurev-virology-100114-055218.

40. Long, S. (2021). SARS-CoV-2 Subgenomic RNAs: Characterization, Utility, and Perspectives. Viruses 13, 1923. 10.3390/v13101923.

41. Mears, H.V., Young, G.R., Sanderson, T., Harvey, R., Crawford, M., Snell, D.M., Fowler, A.S., Hussain, S., Nicod, J., Peacock, T.P., et al. (2022). Emergence of new subgenomic mRNAs in SARS-CoV-2. bioRxiv, 2022.04.20.488895. 10.1101/2022.04.20.488895.

42. Nieto-Torres, J.L., DeDiego, M.L., Verdiá-Báguena, C., Jimenez-Guardeño, J.M., Regla-Nava, J.A., Fernandez-Delgado, R., Castaño-Rodriguez, C., Alcaraz, A., Torres, J., Aguilella, V.M., et al. (2014). Severe Acute Respiratory Syndrome Coronavirus Envelope Protein Ion Channel Activity Promotes Virus Fitness and Pathogenesis. PLoS Pathog. 10, e1004077. 10.1371/journal.ppat.1004077.

43. MisereyLLenkei, S., Trajkovic, K., D’Ambrosio, J.M., Patel, A.J., Čopič, A., Mathur, P., Schauer, K., Goud, B., Albanèse, V., Gautier, R., et al. (2021). A comprehensive library of fluorescent constructs of SARSLCoVL2 proteins and their initial characterisation in different cell types. Biol. Cell 113, 311–328. 10.1111/boc.202000158.

44. Ishida, M., and Bonifacino, J.S. (2019). ARFRP1 functions upstream of ARL1 and ARL5 to coordinate recruitment of distinct tethering factors to the trans-Golgi network. J. Cell Biol. 218, 3681–3696. 10.1083/jcb.201905097.

45. Ma, T., Li, B., Wang, R., Lau, P.K., Huang, Y., Jiang, L., Schekman, R., and Guo, Y. (2018). A mechanism for differential sorting of the planar cell polarity proteins Frizzled6 and Vangl2 at the trans-Golgi network. J. Biol. Chem. 293, 8410–8427. 10.1074/jbc.ra118.001906.

46. Kern, D.M., Sorum, B., Mali, S.S., Hoel, C.M., Sridharan, S., Remis, J.P., Toso, D.B., Kotecha, A., Bautista, D.M., and Brohawn, S.G. (2021). Cryo-EM structure of SARS-CoV-2 ORF3a in lipid nanodiscs. Nat. Struct. Mol. Biol. 28, 573–582. 10.1038/s41594-021-00619-0.

47. Miller, A.N., Houlihan, P.R., Matamala, E., Cabezas-Bratesco, D., Lee, G.Y., Cristofori-Armstrong, B., Dilan, T.L., Sanchez-Martinez, S., Matthies, D., Yan, R., et al. (2023). The SARS-CoV-2 accessory protein Orf3a is not an ion channel, but does interact with trafficking proteins. eLife 12, e84477. 10.7554/elife.84477.

48. Miao, G., Zhao, H., Li, Y., Ji, M., Chen, Y., Shi, Y., Bi, Y., Wang, P., and Zhang, H. (2021). ORF3a of the COVID-19 virus SARS-CoV-2 blocks HOPS complex-mediated assembly of the SNARE complex required for autolysosome formation. Dev. Cell 56, 427–442.e5. 10.1016/j.devcel.2020.12.010.

49. Tseng, Y.-T., Wang, S.-M., Huang, K.-J., and Wang, C.-T. (2014). SARS-CoV envelope protein palmitoylation or nucleocapid association is not required for promoting virus-like particle production. J. Biomed. Sci. 21, 34. 10.1186/1423-0127-21-34.

50. Shepley-McTaggart, A., Sagum, C.A., Oliva, I., Rybakovsky, E., DiGuilio, K., Liang, J., Bedford, M.T., Cassel, J., Sudol, M., Mullin, J.M., et al. (2020). SARS-CoV-2 Envelope (E) Protein Interacts with PDZ-Domain-2 of Host Tight Junction Protein ZO1. bioRxiv, 2020.12.22.422708. 10.1101/2020.12.22.422708.

51. Samavarchi-Tehrani, P., Abdouni, H., Knight, J.D.R., Astori, A., Samson, R., Lin, Z.-Y., Kim, D.-K., Knapp, J.J., St-Germain, J., Go, C.D., et al. (2020). A SARS-CoV-2 – host proximity interactome. bioRxiv, 2020.09.03.282103. 10.1101/2020.09.03.282103.

52. Laurent, E.M.N., Sofianatos, Y., Komarova, A., Gimeno, J.-P., Tehrani, P.S., Kim, D.-K., Abdouni, H., Duhamel, M., Cassonnet, P., Knapp, J.J., et al. (2020). Global BioID-based SARS-CoV-2 proteins proximal interactome unveils novel ties between viral polypeptides and host factors involved in multiple COVID19-associated mechanisms. bioRxiv, 2020.08.28.272955. 10.1101/2020.08.28.272955.

53. May, D.G., Martin-Sancho, L., Anschau, V., Liu, S., Chrisopulos, R.J., Scott, K.L., Halfmann, C.T., Peña, R.D., Pratt, D., Campos, A.R., et al. (2022). A BioID-Derived Proximity Interactome for SARS-CoV-2 Proteins. Viruses 14, 611. 10.3390/v14030611.

54. Jeleń, F., Oleksy, A., Smietana, K., and Otlewski, J. (2003). PDZ domains - common players in the cell signaling. Acta Biochim. Pol. 50, 985–1017. 10.18388/abp.2003_3628.

55. Olmos, Y., Hodgson, L., Mantell, J., Verkade, P., and Carlton, J.G. (2015). ESCRT-III controls nuclear envelope reformation. Nature 522, 236–239. 10.1038/nature14503.

56. Lee, J.K., Jeong, E., Lee, J., Jung, M., Shin, E., Kim, Y., Lee, K., Jung, I., Kim, D., Kim, S., et al. (2018). Directed evolution of CRISPR-Cas9 to increase its specificity. Nat. Commun. 9, 3048. 10.1038/s41467-018-05477-x.

57. Doench, J.G., Fusi, N., Sullender, M., Hegde, M., Vaimberg, E.W., Donovan, K.F., Smith, I., Tothova, Z., Wilen, C., Orchard, R., et al. (2016). Optimized sgRNA design to maximize activity and minimize off-target effects of CRISPR-Cas9. Nat. Biotechnol. 34, 184–191. 10.1038/nbt.3437.

58. Kowarz, E., Löscher, D., and Marschalek, R. (2015). Optimized Sleeping Beauty transposons rapidly generate stable transgenic cell lines. Biotechnol. J. 10, 647–653. 10.1002/biot.201400821.

59. Carlton, J.G., and Martin-Serrano, J. (2007). Parallels Between Cytokinesis and Retroviral Budding: A Role for the ESCRT Machinery. Science 316, 1908–1912. 10.1126/science.1143422.

60. Warren, S.C., Margineanu, A., Alibhai, D., Kelly, D.J., Talbot, C., Alexandrov, Y., Munro, I., Katan, M., Dunsby, C., and French, P.M.W. (2013). Rapid Global Fitting of Large Fluorescence Lifetime Imaging Microscopy Datasets. PLoS ONE 8, e70687. 10.1371/journal.pone.0070687.

61. Cox, J., and Mann, M. (2008). MaxQuant enables high peptide identification rates, individualized p.p.b.-range mass accuracies and proteome-wide protein quantification. Nat. Biotechnol. 26, 1367–1372. 10.1038/nbt.1511.

62. Cox, J., Neuhauser, N., Michalski, A., Scheltema, R.A., Olsen, J.V., and Mann, M. (2011). Andromeda: A Peptide Search Engine Integrated into the MaxQuant Environment. J. Proteome Res. 10, 1794–1805. 10.1021/pr101065j.

63. Tyanova, S., Temu, T., Sinitcyn, P., Carlson, A., Hein, M.Y., Geiger, T., Mann, M., and Cox, J. (2016). The Perseus computational platform for comprehensive analysis of (prote)omics data. Nat. Methods 13, 731–740. 10.1038/nmeth.3901.

64. Mellacheruvu, D., Wright, Z., Couzens, A.L., Lambert, J.-P., St-Denis, N.A., Li, T., Miteva, Y.V., Hauri, S., Sardiu, M.E., Low, T.Y., et al. (2013). The CRAPome: a contaminant repository for affinity purification–mass spectrometry data. Nat. Methods 10, 730–736. 10.1038/nmeth.2557.

65. Cacciabue, M., Currá, A., and Gismondi, M.I. (2019). ViralPlaque: a Fiji macro for automated assessment of viral plaque statistics. PeerJ 7, e7729. 10.7717/peerj.7729.

66. Madeira, F., Park, Y. mi, Lee, J., Buso, N., Gur, T., Madhusoodanan, N., Basutkar, P., Tivey, A.R.N., Potter, S.C., Finn, R.D., et al. (2019). The EMBL-EBI search and sequence analysis tools APIs in 2019. Nucleic Acids Res. 47, W636–W641. 10.1093/nar/gkz268.

67. Waterhouse, A.M., Procter, J.B., Martin, D.M.A., Clamp, M., and Barton, G.J. (2009). Jalview Version 2—a multiple sequence alignment editor and analysis workbench. Bioinformatics 25, 1189–1191. 10.1093/bioinformatics/btp033.

